# Leukemia escapes immunity by imposing a Type-1 regulatory program on neoantigen-specific CD4+ T cells

**DOI:** 10.1101/2024.12.21.629902

**Authors:** Hrishi Venkatesh, Miriam Arroyo, Lynn Heltemes-Harris, Todd P. Knutson, Yinjie Qiu, Allison Haaning, Beau R. Webber, Veronika Bachanova, Michael A. Farrar, Sean I. Tracy

**Affiliations:** Center for Immunology, University of Minnesota, MN, USA; Masonic Cancer Center, University of Minnesota, Minneapolis, MN, USA; Department of Lab Medicine and Pathology, University of Minnesota, Minneapolis, MN, USA; Minnesota Supercomputing Institute, University of Minnesota, MN, USA; Center for Genome Engineering, University of Minnesota, Minneapolis, MN, USA; Stem Cell Institute, University of Minnesota, Minneapolis, MN, USA; Department of Pediatrics, University of Minnesota, Minneapolis, MN, USA; Division of Hematology, Oncology and Transplantation, University of Minnesota, Minneapolis, MN, USA

## Abstract

The significance of endogenous immune surveillance in acute lymphoblastic leukemia (ALL) remains controversial. Using clinical B-ALL samples and a novel mouse model, we show that neoantigen-specific CD4+ T cells are induced to adopt type-1 regulatory (Tr1) function in the leukemia microenvironment. Tr1s then inhibit cytotoxic CD8+ T cells, preventing effective leukemia clearance. Leukemic cells induce Tr1s by phenocopying hematopoietic stem cells, which normally are subject to effective surveillance by this CD4+ subset. This mechanism effectively redirects Tr1 cells from a role in preventing cancer to maladaptively promoting clinical relapse. In mouse models, inhibition of Tr1 expansion with IL10 receptor (IL10R) blockade is insufficient to improve leukemia control. In contrast, combined therapy with a cytotoxic agent and anti-PDL1 blockade eradicated measurable residual disease. This correlates with polarization of the neoantigen-specific CD4+ T-cell population from Tr1 towards Th1 states. Our findings uncover a mechanism that enables leukemic relapse and resolves existing controversies on the role of immune surveillance towards this cancer type. Therapeutic polarization of neoantigen-specific CD4+ T cells towards Th1 states may improve contemporary immune therapies by reshaping the immune microenvironment towards states permissive for cytotoxic attack of residual leukemia.

**Key Points:**

1. B-ALL induces neoantigen-specific CD4+ T cells to adopt Type-1 regulatory states, which protect leukemic cells from immune pressure.
2. Repolarizing neoantigen-specific CD4+ T-cells towards Th1 states eradicates measurable residual disease.

## Introduction

Patients with relapsed B-cell acute lymphoblastic leukemia (B-ALL) continue to experience poor long-term survival. Relapse occurs due to the eventual outgrowth of residual leukemia cells which persist after frontline therapy. Relapse risk is influenced both by the specific molecular and cytogenetic alterations present in leukemic cells at diagnosis and the level of measurable residual disease (MRD) burden after frontline therapy^1,2^. In addition, the anti-leukemia T-cell response may influence relapse risk. This is suggested by findings that higher frequencies of TIM3+ CD4+ T cells at diagnosis correlate with a ∼10-fold higher risk of relapse^3,4^. This increased risk was independent of MRD burden or disease intrinsic factors. Whereas FOXP3+ CD4+ regulatory T cells (Tregs) are broadly known to suppress immune responses, the TIM3+ CD4+ T cells associated with leukemic relapse were reportedly FOXP3-negative. We previously found that TIM3+/FOXP3-CD4+ T cells did not harbor hallmarks of functional deficits, including exhaustion^5^. The capacity for FOXP3-CD4+ T-cell subsets to actively suppress anti-leukemia immune responses has not been well described. Therefore, explanations for the observed association of relapse risk with endogenous TIM3+/FOXP3-CD4+ T cells remain largely unknown.

Here, we demonstrate that B-ALL induces leukemia-specific CD4+ T-cells to adopt Type-1 regulatory (Tr1) states. Tr1s are well known for inducing peripheral tolerance by delivery of immunosuppressive signals. We show that Tr1s are the predominant population that comprises the TIM3+/FOXP3-CD4+ subset, previously shown to be associated with relapse. Remarkably, B-ALL clearance with PD-L1 blockade and tyrosine kinase inhibitor therapy is associated with rescue of leukemia-specific CD4 T cells from Tr1 states. Our investigations suggest that effective anti-leukemia immune prevails when leukemia-specific CD4+ T-cells are converted from Tr1 to Th1 functions.

## Methods

### Mice

CD45.1^+^ and CD45.2^+^ C57BL/6 mice were acquired from Charles River Laboratories. B6 Thy1.1 mice were acquired from Jackson laboratories. IL10-GFP reporter mice that have been described before^6^ were a gift from Dr. Sara Hamilton-Hart at the University of Minnesota. SMARTA TCR transgenic mice that have been described before^7^ were a gift from Dr. Dave Masopust at the University of Minnesota. Mice were housed at the University of Minnesota in specific pathogen-free conditions, and all experiments were approved by the Institutional Animal Care and Use Committee. Mice were generally 8-10 weeks old, but ranged 7–16 weeks. Mice were randomly selected for experiments in age-matched cohorts.

### Leukemia Model

The murine BCR-ABL^+^ leukemia cell line (LM138) was generated as previously described^8,9^. Briefly, the cell line was generated by transducing the bone marrow of Cdkn2a^−/−^ mice (that specifically lack but not *Ink4a*) mice with viral supernatant containing a BCR-ABL (p190)-IRES-GFP retrovirus. The cells were maintained for adoptive transfer as described before^8,9^. Additional leukemia cell lines used include a cell line generated spontaneously in *Pax5^+/−^ x Ebf1^+/−^* mice.

### Making HV1 TCR KI Mice

The HV1 TCR KI mice were made using the approach described recently by the Stromnes and Webber lab^10^. Briefly, the TCRα and TCR; sequences from the Dominant LM138 leukemia-specific T cell clone described in our previous Blood paper was synthesized into a gene block, and were knocked into the rAAV-mTRAC-GSG-T2A vector using the Gateway LR Clonase II Enzyme Mix (ThermoFisher Scientific, Waltham, MA). pAAV constructs were sent to SignaGen Laboratories for commercial rAAV production. Zygotes from superovulated females were electroporated with a Cas9-CRISPR RNP complex containing a guide RNA targeting exon 1 of the TRAC locus (UUCUGGGUUCUGGAUGUCUG), as well as 2uL of the rAAV containing the dominant clonotype TCR (1.5 × 10^9^ GC/μl). The electroporated zygotes were injected into pseudo-pregnant CD45.2 mice to generate the TCR KI mice. The TCR knock-in was validated using flow cytometry staining for the TCR; chain (TCR V;7), and by qPCR for the dominant clonotype TCR, as described before^10^. The primers used were *WT Fwd*: CTCTGGTGTGAGTGCTATTC, *TCR KI FWD*: ATTTCTCCTCTACCGCCACA and *UNIV JXN REV*: CAAGAGAAGACAGGAAGGTGAG.

### T cell isolation from leukemic mice

CD45.2^+^ C57BL/6 Mice were challenged with 2500 LM138 cells via tail vein injection. The mice were sacrificed at the indicated time-points post-leukemia challenge. The spleen and bone marrow were isolated. The spleen was processed into a single-cell suspension using frosted slides while the bone marrow was flushed using an insulin syringe. The single cell suspensions from the spleen and bone marrow were filtered using a 70µM filter (BD Biosciences). LM138 cells (or other leukemia cell lines) from the spleen were depleted by staining with biotinylated antibodies against CD19 and B220, followed by depletion using Streptavidin Rapidpheres (StemCell). T cells from the LM138-depleted spleen and bone marrow were isolated using the EasySep CD90.2 positive-selection kit (StemCell). HV1 TCR knock-in CD4 T cells were enriched by staining for either TCR V;7 (the TCR; chain expressed by HV1 T cells) or CD45.2, followed by enrichment using the EasySep PE Selection Kit or EasySep APC Selection Kit (StemCell).

### Ex-vivo Restimulation with PMA and Ionomycin

The isolated enriched murine T cells were incubated with PMA (20ng/mL) and Ionomycin (1µg/mL), or DMSO, in a 96-well plate. 1 hour later, 1µg/mL GolgiPlug and 1.5µg/mL GolgiStop (BD BioSciences) were added. 5 hours after the addition of GolgiPlug and GolgiStop, the cells were isolated for flow cytometry.

### In-vitro peptide screen

CD4 T cells from HV1 TCR KI mice were enriched using by staining with a biotin antibody cocktail (CD8, CD11b, CD11c, CD19, B220, F4/80, GR1, NK1.1, Ter119, γδTCR), followed by negative selection using Streptavidin Microbeads and LD columns (Miltenyi Biotec), per the manufacturer’s instructions. Splenic APCs were isolated via depletion of CD90.2^+^ cells from the spleens of C57BL/6 mice using Streptavidin Microbeads and LS columns (Miltenyi Biotec). HV1 T cells and splenic APCs were cultured in 96-well plates (5e4 HV1 T cells and 2e4 splenic APCs per well) in 200uL complete RPMI. Each peptide in the BCR-ABL peptide library was reconstituted in DMSO (Fischer) at 20mg/mL and added to one well of a 96-well plate at a concentration of 100µg/mL. As a positive control, anti-CD3 and anti-CD28 antibodies were added at 2µg/mL. After 72hrs, activation markers (CD25, CD44, PD1) on the HV1 T cells were analyzed by flow cytometry.

### Flow cytometry

Flow cytometry antibodies are in Table 1, below. For flow cytometric analyses and FACS sorts, T cell-enriched or unfractionated spleen cells were stained in PBS + 2% FBS + 2mM EDTA + 0.05% Azide for 20 min with corresponding antibodies and washed. Intracellular staining was done using the FOXP3-transcription factor staining kit (Invitrogen), as per the manufacturer’s instructions. Cells from enriched fractions were analyzed on LSRII or Fortessa cytometers (BD Biosciences, San Jose, CA), or on Cytek Aurora cytometers (Cytek Biosciences, Fremont, CA) and data were analyzed in FlowJo (TreeStar, Ashland, OR).

**Table 1:**

| ANTIBODY | SOURCE | IDENTIFIER |
| --- | --- | --- |
| Anti-mouse CD4 eF450 | eBioScience | 48-0041-82 |
| Anti-mouse CD4 PE | eBioScience | 12-0042-82 |
| Anti-mouse CD4 BUV496 | BD Bioscience | 612952 |
| Anti-mouse CD8 BV605 | Biolegend | 100744 |
| Anti-mouse CD8 BV650 | BD Bioscience | 563234 |
| Anti-mouse CD8b Biotin | Biolegend | 126604 |
| Anti-mouse CD11b Biotin | Biolegend | 101204 |
| Anti-mouse CD11c Biotin | eBioScience | 13-0114-85 |
| Anti-mouse CD11c FITC | eBioScience | 11-0114-85 |
| Anti-mouse MHCII (I-A) PE | eBioScience | 12-5322-82 |
| Anti-mouse MHCII (I-A/I-E) eF450 | eBioScience | 48-5321-82 |
| Anti-mouse CD19 Biotin | Biolegend | 115504 |
| Anti-mouse CD19 BV421 | Biolegend | 115538 |
| Anti-mouse CD19 BV605 | BD Bioscience | 563148 |
| Anti-mouse CD25 PE | eBioScience | 12-0251-83 |
| Anti-mouse CD25 PerCP-Cy5.5 | Biolegend | 102030 |
| Anti-mouse/human CD44 BV786 | Biolegend | 103059 |
| Anti-mouse/human CD44 PerCP-Cy5.5 | Biolegend | 103032 |
| Anti-mouse CD45.1 FITC | eBioScience | 11-0453-85 |
| Anti-mouse CD45.1 PE | Biolegend | 110707 |
| Anti-mouse CD45.2 APC | Biolegend | 109814 |
| Anti-mouse CD45R(B220) Biotin | Biolegend | 103204 |
| Anti-mouse CD90.1 PE | Biolegend | 202524 |
| Anti-mouse CD90.1 PE-Cy7 | eBioScience | 25-0900-82 |
| Anti-mouse CD90.1 BUV737 | BD Bioscience | 612837 |
| Anti-mouse CD90.2 BV786 | Biolegend | 564365 |
| Anti-mouse CD90.2 BUV395 | BD Bioscience | 565257 |
| Anti-mouse F4/80 Biotin | Biolegend | 123106 |
| Anti-mouse Gr1 Biotin | Biolegend | 108404 |
| Anti-mouse NK1.1 Biotin | Biolegend | 108704 |
| Anti-mouse Ter119 Biotin | Biolegend | 116204 |
| Anti-mouse $\gamma\delta$TCR Biotin | Biolegend | 118103 |
| Anti-mouse TCR Vb7 PE | Biolegend | 118307 |
| Anti-mouse TCR Vb7 FITC | Biolegend | 118305 |
| Ghost Red Viability Dye | Tonbo | 50-105-2989 |
| Anti-mouse PD1 PE-Cy7 | Biolegend | 109109 |
| Anti-mouse PD1 BV711 | Biolegend | 135231 |
| Anti-mouse Tim3 BV605 | Biolegend | 119721 |
| Anti-mouse Lag3 PerCP-Cy5.5 | Biolegend | 125212 |
| Anti-mouse CXCR5 BV650 | Biolegend | 145517 |
| Anti-mouse CD40L APC | Biolegend | 157009 |
| Anti-mouse IFNγ APC | Biolegend | 505810 |
| Anti-mouse TNF BV421 | BD Bioscience | 563387 |
| Anti-mouse IL10 FITC | Biolegend | 505005 |
| Anti-mouse IL10 PE | Biolegend | 505007 |
| Anti-mouse EOMES PE-Cy7 | eBioScience | 25-4875-82 |
| Anti-mouse FOXP3 AF488 | eBioScience | 53-5773-82 |
| Anti-mouse FOXP3 eF450 | eBioScience | 48-5773-82 |
| Anti-mouse FOXP3 PerCP-Cy5.5 | eBioScience | 45-5773-82 |

### In-Vitro Suppression Assay

CD4 T cells were enriched from SMARTA TCR transgenic mice by staining with a biotin antibody cocktail (CD8, CD11b, CD11c, CD19, B220, F4/80, GR1, NK1.1, Ter119, γδTCR), followed by negative selection using Streptavidin Microbeads and LD columns (Miltenyi Biotec), per the manufacturer’s instructions. HSPCs were enriched from the bone marrow using a biotin depletion cocktail (CD4+, CD8, CD11b, GR1, Ter119) and Streptavidin Rapidspheres (StemCell). Splenic APCs were isolated via depletion of CD90.2+ cells from the spleens of C57BL/6 mice using Streptavidin Microbeads and LS columns (Miltenyi Biotec). SMARTA T cells and the indicated APCs were cultured in 6 well plates in complete RPMI at a 2.5:1 ratio and a density of 0.5× 10^6^ SMARTA T cells/mL for 72hrs with 50µg/mL Gp66 peptide (stock: 20mg/mL in DMSO). 3 days later, the SMARTA T cells and Tregs from a FOXP3-GFP mouse were sorted using a BD FACSAria sorter. 5e4 SMARTA T cells or Tregs were cultured with 1e6 CTV-labeled T cells from a CD45.1+ mouse, as well as 1e6 T cell-depleted splenocytes from a CD45.2+ C57BL/6 mouse with 1µg/mL soluble anti-CD3 antibody (Clone 17A2, Tonbo Biosciences) in a 96-well plate. Cells were analyzed by flow cytometry and proliferation of bystander cells was assessed.

### Ex-vivo Suppression Assay

HV1-T cells were enriched from leukemic mice at the indicated time points as indicated before, and then sorted using a BD FACSAria sorter. FOXP3-GFP^+^ Tregs and FOXP-GFP^−^ T conv CD4+-T cells were also sorted from a WT FOXP3-GFP mouse. 5e4 HV1 T cells, Tconv CD4s or Tregs were cultured with 1e6 CTV-labeled T cells from a CD45.1+ mouse, as well as 1e6 T cell-depleted splenocytes from a CD45.2+ C57BL/6 mouse with 1µg/mL soluble anti-CD3 antibody (Clone 17A2, BioXCell) in a 96-well plate. Cells were analyzed by flow cytometry and proliferation of bystander cells was assessed.

### Nilotinib and/or blocking antibody treatments

Mice challenged with 2500 LM138 leukemia cells were bled at day 13 post-leukemia challenge to confirm leukemia engraftment. Mice were treated 75mg/kg Nilotinib (MedChemExpress) on days 14-18 via oral gavage, as well as anti-mouse PDL1 (clone 10F.9G2) or anti-mouse IL10R (clone 1B1.3A) or isotype control antibodies (BioXCell) at 10mg/kg on days 14, 16 and 18 via intraperitoneal injection.

### Single-cell ATAC sequencing of HV1 CD4 T cells

For scATAC-seq, we sorted HV1s from the spleens from naïve HV1-TCR knock-in mice, and mice challenged with LM138 leukemia cells at the indicated time points post leukemia challenge (100,000 HV1s per group). Sorted cells were centrifuged at 300 × g for 5 min at 4 °C. Cells were resuspended in 100 µl chilled lysis buffer, which consists of 10 mM Tris-HCl. pH 7.4 (T2914, Sigma-Aldrich), 10 mM NaCl (S5150, Sigma), 3 mM MgCl2 (M1028, Sigma-Aldrich), 0.1% Tween 20 (BP337, Thermo Fisher Scientific), 0.1% IGEPAL CA-630 (I8896, Sigma-Aldrich), 0.01% digitonin (BN2006, Thermo Fisher Scientific), 1% BSA (130-091-376, Miltenyi Biotec) and nuclease-free water, and incubated on ice for 5 min. After incubation, 1 ml chilled wash buffer, including 10 mM Tris-HCl, pH 7.4 (T2914, Sigma-Aldrich), 10 mM NaCl (S5150, Sigma-Aldrich), 3 mM MgCl2 (M1028, Sigma-Aldrich), 0.1% Tween 20 (BP337, Thermo Fisher Scientific), 1% BSA (130-091-376, Miltenyi Biotec) and nuclease-free water, was added to lysed cells and centrifuged at 500 × g for 5 min at 4 °C. Cells were resuspended in diluted nuclei buffer, and nuclei permeabilization was determined using a Countess II (Thermo Fisher Scientific). Cells were captured on the 10x Chromium, and library preparation was generated using the Chromium Next GEM Single Cell ATAC library kit v1.1 (PN-1000164, 10x Genomics), following the manufacturer’s recommendations. Post-library quality control was performed for sizing and quantification using the Agilent Bioanalyzer (Agilent 2100 Bioanalyzer, Agilent Technologies) and KAPA library quantification kit for Illumina platforms (KR0405, Roche). All libraries were sequenced on a NovaSeq 6000 with 2 × 150 bp paired-end reads (Illumina).

### Analysis of scATAC-seq data

Single-cell ATAC seq data was processed using cellranger-atac (V 2.1.0) for each time point of the study separately using mouse refdata-cellranger-arc-mm10-2020-A-2.0.0 as the reference. Fragments.tsv.gz and singlecell.csv output files for each time point were aggregated with depth normalization using cellranger-atac aggr function. To analyze the dataset, Signac (V 1.9.0) was built under R (V 4.0.0). Data was filtered using recommended parameters peak_region_fragments > 3000, peak_region_fragments < 100000, pct_reads_in_peaks > 40, blacklist_ratio < 0.025, nucleosome_signal < 4, TSS.enrichment > 2 options and the resolution was set to 0.5 for the rest analysis. Gene activity was predicted using GeneActivity function and the cluster identity was determined by examining the activities of known marker genes. FindMarkers was used to identify top differential accessible peaks (min.pct = 0.05) and CoveragePlot was used to visualize the chromatin accessibility data. The data was deposited in the Gene Expression Omnibus database (accession number: GSE***).

### Single-cell RNA/TCR sequencing of HV1 CD4 T cells and CD8 T cells

10,000 CD45.2^+^ HV1-CD4s were transferred into congenic CD45.1^+^ mice, followed by a challenge with 2500 LM138 leukemia cells the next day. The leukemic CD45.1^+^ mice were either not treated, or treated at day 14 post-challenge with nilotinib and isotype control/anti-PDL1 blockade/anti-IL10R blockade as described above. Single cell suspensions were prepared from the spleens of the untreated leukemic CD45.1^+^ mice at day 13 post-leukemia challenge, as well as from leukemic CD45.1^+^ mice treated with nilotinib + isotype, IL10R, or PDL1 blocking antibodies at day 21 post-leukemia challenge. Cells from each treatment group were stained using unique TotalSeq hashtag oligonucleotide (HTO) antibodies (BioLegend, cat. C0301-C0305). Cell surface protein levels were measured for six markers: CD4, CD39, CD49b, PD1, TIM3, and LAG3 using TotalSeq CITE-seq antibody derived tags (ADT) (BioLegend cat. C0001, C0834, C0421, C0004, C0003, C0378). An equal number of CD45.2^+^ HV1 CD4^+^ T cells and CD44^+^ CD45.1^+^ CD8 T cells were sorted using a BD FACSAria sorter (25,000 cells per treatment arm). In addition, 5000 naïve HV1-CD4^+^ T cells were sorted from naïve HV1 TCR-knock-in mice. The sorted T cells were resuspended at 10^6 ml in 50% FBS in 1× PBS before being counted and captured using 10x Genomics Single Cell 5’ Solution. Cells were processed for mRNA and TCR expression levels using the Chromium Next GEM Single Cell 5’ Reagent Kit v2 (10x Genomics). Two libraries, gene expression (GEX) and feature barcode (HTOs and CITE-seq ADTs), were pooled and sequenced in one lane of an Illumina NovaSeq 6000 instrument, using 2×151 bp paired-end read chemistry. The TCR library was sequenced on the same instrument in a different lane.

### Analysis of scRNA-seq data

Illumina indexes were used to demultiplex the gene expression and feature barcode libraries into sets of fastq files. Unique 10X Genomics barcodes were recovered from individual gel bead in emulsion (GEM) droplets and these barcodes link sequencing results from the GEX, ADT, HTO, and TCR libraries. The GEX reads were pseudoaligned against the mouse transcriptome (mm10) using a kallisto (v. 0.48.0) and bustools (v. 0.39.3) wrapper pipeline, kb, using python (v. 3.7.10). The 10X Genomics barcode allow-list, “737K-august-2016” was used to limit and correct barcodes with a Hamming distance of 1 from any known barcode. The feature barcode library was processed using kite (https://github.com/pachterlab/kite; commit aafc8c8), which utilizes kallisto and bustools functions. An HTO and ADT oligonucleotide allow-list was generated and a kallisto index was generated. The number of features barcodes were counted for each GEM. Raw TCR library fastq files were mapped and assembled into contigs using the 10X Genomics reference genome (refdata-cellranger-vdj-GRCm38-alts-ensembl-5.0.0) and GEM-specific TCR clonotyping was performed using the cellranger vdj function. GEMs were classified as cell-singlets, multiplets, or negatives based on the expression of normalized (centered log ratio) hashtag counts using GMM-demux (v. 0.2.1.3) with default settings. Raw CITE-seq ADT counts were normalized using the dsb R package (v. 1.0.2), which uses non-cell containing GEMs to determine background levels of ADT abundance. Count tables were imported into R (v. 4.2.0) (https://www.R-project.org/) and processed with functions from the tidyverse package (v. 1.3.2). The dataset was filtered to include only GEM barcodes classified as cell singlets (based on HTO abundance) and further processed using the Seurat R package (ver. 4.2.0). Gene expression data were normalized using the SCTransform method and GEMs were evaluated using the Seurat CellCycleScoring function. 3,000 variable genes were discovered using the vst algorithm. The CITE-seq ADT abundance levels were used to classify GEMs based on CD4 expression, and the dataset was subset to include only CD4 GEMs in downstream analyses. The CD4 set was subjected to PCA and UMAP dimensionality reduction. We explored a variety of clustering resolutions using the default Seurat clustering algorithm (Louvain) in PCA space and chose the 0.5 resolution based on known differences in marker gene expression. The Seurat FindMarkers function (using the Wilcoxon rank sum test) was used to identify differentially expressed genes between clusters or samples/clusters. Plots were generated using the ggplot2 (ver. 3.4.0) R package with the viridis R package (ver. 0.6.2) color scale.

Gene set enrichment analysis (GSEA) was performed using the clusterProfiler R package (ver. 4.4.4) with the GSEA function. The msigdbr (v. 7.5.1) R package provided homologous mouse gene sets (derived from the human MSigDB gene sets). We combined these lists with custom mouse gene sets extracted from recent T cell publications and performed the analysis with default options. The gene expression dataset was preranked based on the log2 fold change of genes expressed in between comparison groups.

Single cell gene expression, feature barcode (CITE-seq ADT and HTO), and TCR locus sequencing data were deposited in the Gene Expression Omnibus database (accession number: GSE***, GSE***).

### Differential expression and gene set enrichment analysis of Tr1s and Thsc from previously published datasets

Raw gene counts from the recently-published bulk RNA-seq dataset comparing OTII-CD4^+^ T cells activated by conventional dendritic cells (T_DC_) and hematopoietic stem cells (T_HSC_)^11^ were used with DESeq2 (v1.42.1) to identify genes differentially expressed between the two cell types (see Supplementary Table 2). CD4^+^ T cells from mice with a live leukemia challenge and mice with heat-killed leukemia immunization were subsetted from our previously-published scRNA-seq dataset^5^ using the Seurat R package (ver 4.2). Differentially-expressed genes between Tr1-like CD4^+^ T cells from mice with a live leukemia challenge over CD4^+^ T cells from mice with heat-killed leukemia were identified using the Seurat R package (ver 4.2) as described above (see Supplementary Table 3). Genes upregulated or downregulated in T_HSC,_ as well as the genes associated with a previously published co-inhibitory module were included as gene sets for gene set enrichment analysis (GSEA) using the R package clusterProfiler (ver 4.10.1), as described above.

### Analysis of TARGET Acute Lymphoblastic Leukemia Database using UCSC Xena

The results published here are in whole or part based upon data generated by the Therapeutically Applicable Research to Generate Effective Treatments (https://www.cancer.gov/ccg/research/genome-sequencing/target) initiative, phs000218. Data used for this analysis are available at the Genomic Data Commons (https://portal.gdc.cancer.gov). The TARGET-ALL-P2 cohort (dbGAP sub-study ID phs000464) is comprised of high-risk pediatric patients with precursor B-cell ALL, most of whom experienced an early bone marrow relapse (within 4 years of initial diagnosis). Tissues were collected from patients enrolled in COG biology studies and clinical trials. mRNA analysis was performed by GDC protocols (https://docs.gdc.cancer.gov/Data/Bioinformatics_Pipelines/Expression_mRNA_Pipeline/), and quantified as upper quartile normalized FPKM (FPKM-UQ) and aligned to reads from the GRCh38 reference genome. The TARGET-ALL-P1 cohort **(dbGAP sub-study ID** phs000463) is comprised of 207 samples from newly diagnosed pediatric patients diagnosed between March 2000 and April 2003 that were defined as at high risk for relapse, and were treated on the POG9906 clinical trial. RNA expression was quantified by Human Genome U133 Plus 2.0 Arrays (Affymetrix). Details of RNA quantification for both cohorts have been published previously (citations: **19470474,** 21360654, 22897847). A Tr1 gene signature comprised of the 241 DEGs found to be upregulated in Tr1 versus Th1 cells (using a -logFDR of 2.0); see **Supplementary Table 1**. Kaplan-Meier survival analysis with log-rank testing for overall survival was conducted using the UCSC Xena platform ((32444850) to assess the association of the 241-gene Tr1 signature, with equal weighting of each gene. Analysis was repeated using a 19-gene Tr1 signature was comprised of genes known to be important in Tr1 induction, maintenance, and effector function.

### Statistical Analysis

Comparisons of two groups were done by either paired t-test (paired, normal data), Wilcoxon matched-pairs test (paired, non-normal data), t-test (non-paired, normal data) or Mann–Whitney (non-paired, non-normal data); tests were always two-sided. Comparison of three or more groups was done by one-way ANOVA (non-paired, normal data) or Kruskal–Wallis (non-paired, non-normal data). Analysis of survival was done using the log-rank test. P < 0.05 was considered significant. Statistics were calculated using Prism (GraphPad Software). All data, except those specifically mentioned in Figure legends, are displayed as mean ± s.d.

### Data and code availability

scRNA-seq and scATAC-seq data have been deposited to the Gene Expression Omnibus (GEO) and are publicly available based on GEO accession numbers. The data for the scRNA-seq and scATAC-seq analysis has been deposited on Zenodo.

### Materials availability

The HV1 TCR knock-in mice, as well as the LM138 and Pax5^+/−^ x Ebf1^+/−^ leukemia cell lines are available on reasonable request from the lead contact, Dr. Sean I Tracy.

## Results

### Human bone marrow TIM3+ CD4+ T cells from patients with newly diagnosed B-ALL are predominantly Tr1s, and Tr1 gene signatures correlate with inferior survival

To characterize the transcriptome of relapse-associated TIM3+ CD4+ T cells, we re-analyzed our previously published scRNA-seq dataset of T cells present in diagnostic bone marrow biopsies of patients with Ph+ B-ALL^5^. We compared differentially-expressed genes (DEGs) between three clusters of identified CD4+ T cells (**Figure 1A**). One cluster (“Th1s”) demonstrated partial downregulation of *TCF7* and *IL7R,* and upregulation of *TNF*, *CD40L*, and *AHNAK*, which encodes a calcium channel scaffolding protein that is critical for effector/memory CD4+ T-cell activation^12^. Compared to Th1s, a second cluster cells revealed 347 differentially overexpressed genes, of which 246 genes were overexpressed (**Figure 1B, C**). These included the inhibitory receptors *PDCD1, HAVCR2, CTLA4, TIGIT,* and *LAG3.* The specific expression of *HAVCR2* (which encodes for TIM3) was limited to this cluster, suggesting these cells are representative of the TIM3+ CD4+ T cells previously reported to be associated with relapse^3,4^. This cluster also overexpressed *IL10, PRDM1,* and *CHI3L2.* Additional upregulated genes included those with cytotoxic function, such as *NKG7, GZMB, GZMH, GZMK, PRF1*, and several genes related to the CCR5 chemokine axis including *CCL3, CCL4,* and *CCL5.* The TIM3+ cluster also overexpressed *Mki67*, implying active proliferation. *FOXP3* was not expressed. Overall, this transcriptional profile is consistent with a Type-1 regulatory (Tr1) state^13–15^. Next, we treated the 246 upregulated genes in the Tr1 cluster as an equal-weighted gene signature (**Supplementary Table 1**), and examined impact on clinical outcomes in patients with B-ALL. The Tr1 signature was associated with inferior overall survival among a cohort of high-risk pediatric patients with B-ALL (TARGET-ALL-P2) (**Figure 1D-F**). We then derived a more limited 19-gene signature comprised of Tr1-associated genes from literature review^15^. The 19-gene signature was associated with inferior overall survival in the TARGET-ALL-P2 panel as well as a separate pediatric cohort (TARGET-ALL-P1). This data suggests that presence of TIM3+/FOXP3-CD4+ T cells with a Tr1 transcriptional signature at diagnosis, which yield poor prognosis in B-ALL.

**Figure 1.**
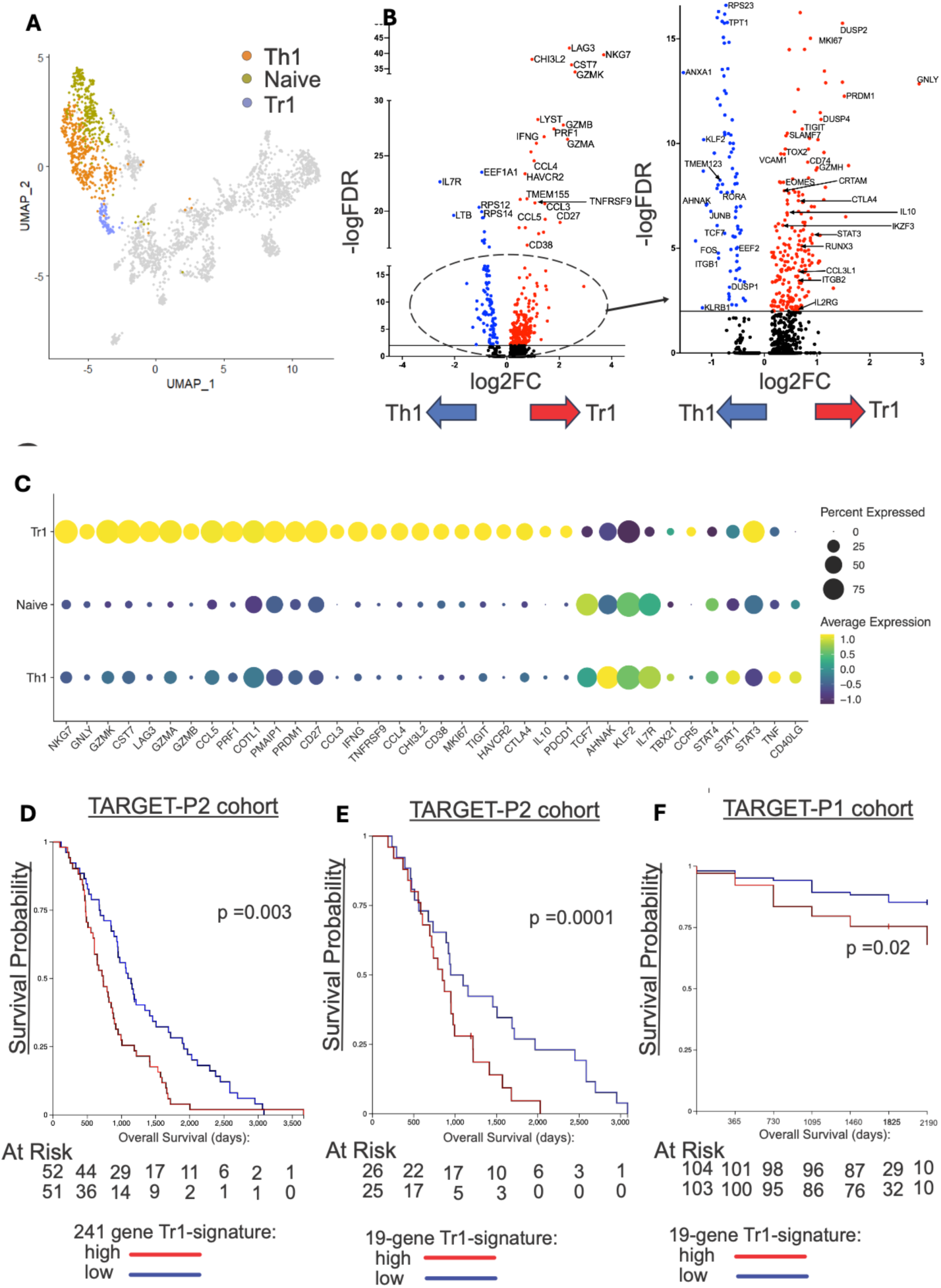
Type-1 regulatory signatures correlate with inferior overall survival among pediatric patients with B-ALL. A) scRNA-seq of diagnostic bone marrow aspirates (n = 5) of adult patients with B-ALL. Pooled CD4+ T cells with annotated differentiation state highlighted; Th1 = 394, Tr1 = 69, Naïve = 270. (B) Volcano plots of differentially regulated genes between Tr1 and Th1 cells. Horizontal line indicates a -logFDR value of 2.0. (C) Dot plots of select genes from CD4+ T-cell states. (D) Kaplan-Meier curves of the GDC-TARGET-ALL-P2 cohort as stratified into high- and low-risk groups by expression of a gene signature comprised of DEGs (n = 241) upregulated in Tr1 compared to Th1 cells. (E) As in D, but applying a gene signature (n = 19) derived from canonical Tr1 genes; patients were stratified by highest versus lowest quartile of expression. (F) Kaplan-Meier curves for a separate cohort of high-risk pediatric patients and stratified by the 19-gene Tr1 gene signature.

### TIM3^+^ CD4+ T cells in mouse models of B-ALL are predominantly Tr1s

We previously reported that TIM3+/FOXP3-CD4+ T cells spontaneously arise in a mouse model of BCR-ABL+ B-ALL. To understand if this subset was similar to the Tr1s observed in clinical specimens, we examined DEGs that were overexpressed in this subset compared to a control population of FOXP3-CD4+ T cells. The TIM3+/FOXP3-CD4+ T cells had upregulated *Il10, Lag3,* and *Maf,* consistent with a Tr1 state (**Figure 2A**). To validate a Tr1 identify, we challenged IL10-GFP reporter mice with the murine BCR-ABL+ leukemia cell line, LM138^8,9^, and analyzed CD4+ T cells at post-leukemia challenge timepoints. We observed a progressive increase in expression of IL10, TIM3, and LAG3 among FOXP3-, antigen-experienced (CD44+/ PD1+) CD4+ T cells during leukemia outgrowth (**Figure 2B, C**). We observed similar production of IL10 in TIM3+/LAG3+ CD4+ T cells from mice with *Pax5^+/−^ x Ebf^+/−^* leukemia, demonstrating that this phenotype is also generalizable to BCR-ABL-B-ALL (**Supplementary Figure 1**). Thus, both BCR-ABL+ and BCR-ABL-murine B-ALL induce CD4+ T cells that phenotypically resemble Tr1s.

**Figure 2:**
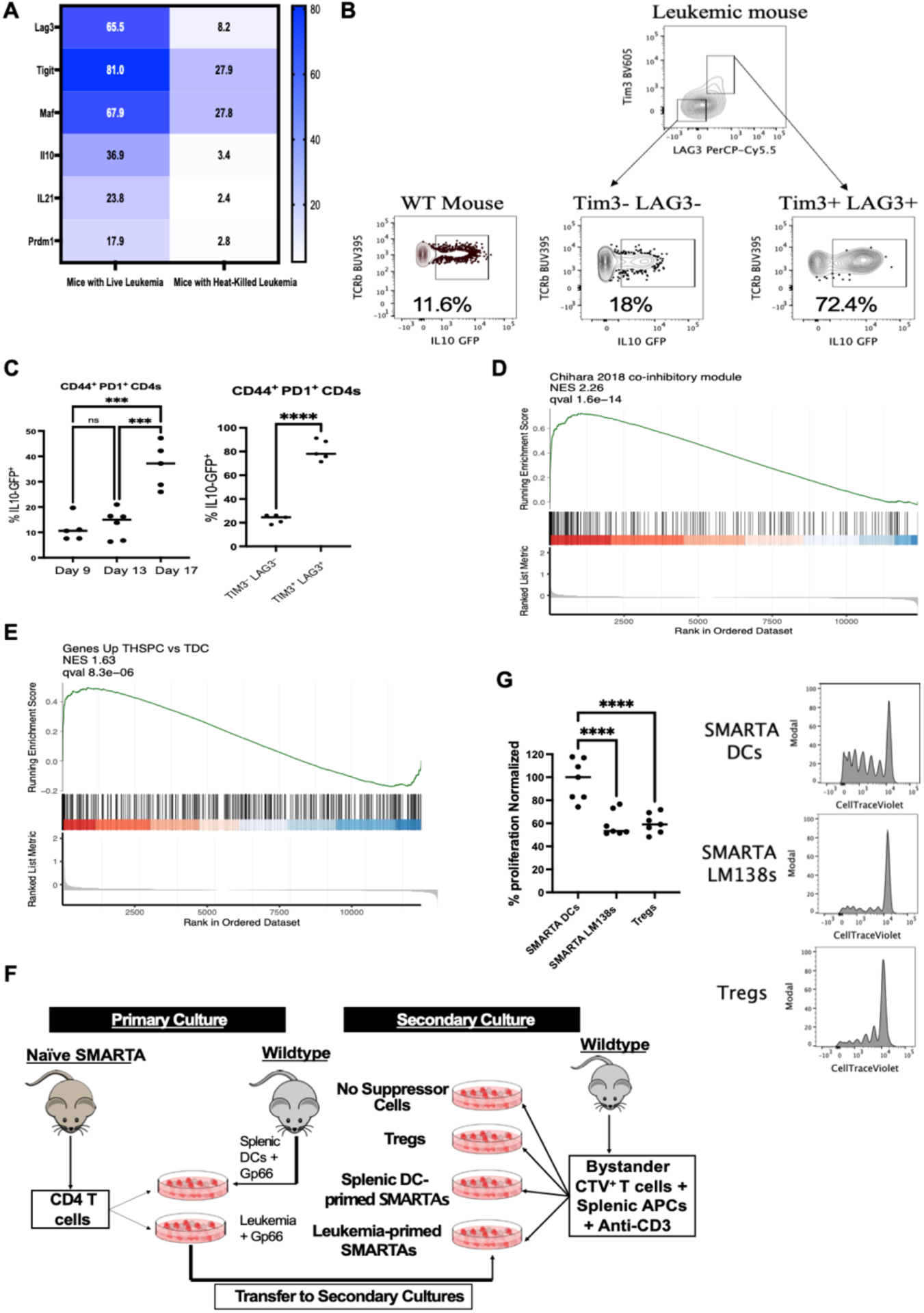
B-ALL induces dysfunctional T cells with a Tr1 phenotype that resemble HSC-induced suppressive CD4+ T cells. A) Heatmap showing the proportion of cells that express Tr1-associated genes in a previously-identified cluster of “helper-cytotoxic-exhausted” CD4+ T cells that specifically arise during leukemia, compared to FOXP3-CD4+ T cells from mice vaccinated with heat-killed leukemia. B,C) IL10-GFP reporter mice were challenged with 2500 LM138 leukemia cells. Representative flow plots and graphs showing the upregulation of IL10 in CD4+ T cells from leukemic mice, stratified by co-expression of TIM3 and LAG3. The results are a summary of 2 experiments. D) GSEA plots showing the enrichment of genes associated with a co-inhibitory gene module from Chihara et al. within the Tr1 CD4+ T-cell cluster from untreated leukemic mice. E) GSEA plots showing the enrichment of genes upregulated in HSC-primed OTII T cells from Hernández-Malmierca et al.^11^, within the Tr1 CD4+ T-cell cluster from untreated leukemic mice. F) SMARTA T cells were activated with cognate peptide for 3 days using splenic DCs or LM138 leukemia cells. Activated SMARTA T cells were sorted and cultured with bystander CTV+ T cells with APCs and anti-CD3. G) Graphs and representative plots show proportion of bystander T cells that proliferated. The results are a summary of 2 experiments. Individual values are shown in C and F. Statistical significance was determined using a one-way ANOVA, followed by the Holm-Sidak’s multiple comparisons test. ***p<0.001, ****p<0.0001, ns: not significant.

To determine whether this CD4 subset can arise from priming by B-ALL cells, we co-cultured SMARTA transgenic CD4+ T cells with LM138 cells pulsed with their cognate peptide Gp66^7^. After 72 hours of co-culture, primed SMARTA T cells were transferred to secondary cultures with bystander T cells. GSEA was performed. Primed CD4+ T cells were enriched with a co-inhibitory gene module (**Figure 2D**)^16^. They were also enriched for a gene module expressed by CD4+ OTII T cells that had adopted a Tr1 state after being primed by hematopoietic stem cells (HSCs) (**Figure 2D, 2E**)^11^. As expected, LM138-primed SMARTAs remained FOXP3 negative but suppressed bystander T cells.

The degree of suppression was comparable to that of purified FOXP3+ regulatory T cells (**Figure 2F, G**). Thus, leukemic cells prime CD4+ T cells towards a FOXP3-suppressive state via presentation of cognate antigen, similar to the actions of HSCs.

### Characterization of a leukemia neoantigen-specific TCR knock-in mouse

Our prior work examined polyclonal CD4+ T cells from wild-type B6 mice challenged with leukemia^5^. This approach could not discriminate between leukemia-specific CD4+ T cells and bystander clones, or recently activated T cells from chronically-stimulated ones. To overcome these limitations, we developed a T-cell receptor (TCR) knock-in mouse model that produces leukemia-specific CD4+ T cells. We had previously observed that CD4+ T cells clonally expanded in LM138 leukemia-bearing mice during dual therapy with the BCR-ABL inhibitor nilotinib and anti-PDL1 blockade (**Figure 3A**)^5^. In those studies, one dominant clonotype of CD4+ T cells emerged. We suspected this clone specifically recognized an unknown leukemia antigen. We sequenced the *Tcra* and *Tcrb* genes of this dominant clonotype and used CRISPR/CAS9-based engineering to knock-in both TCR genes into the *Trac* locus (**Figure 3B, C**)^17^. T cells in this mouse developed normally and expressed the expected TCR V*β*7chain (**Figure 3D**). Subsequent PCR analysis confirmed that the knock-in T cells expressed the expected *Tcra (TRAV13-1)* and *Tcrb* (*TRBV29*) genes (not shown). We hypothesized that HV1 CD4+ T cells may recognize a neoantigenic region of the BCR-ABL fusion oncogene. Multiple candidate regions were considered. The fusion region between BCR and ABL forms a well-characterized neoantigen^9,18,19^. Furthermore, the LM138 cell line expresses a human-encoded version of BCR-ABL, which is highly conserved but not identical to murine BCR and ABL; thus, non-conserved amino acid residues may function as neoantigens in a murine background. Finally, LM138 cells spontaneously acquire mutations in the BCR-ABL tyrosine kinase domain during treatment with tyrosine kinase inhibitors, which may create neoantigens. To identify the actual epitope recognized by HV1 cells, we designed a peptide library that spanned the entire human BCR-ABL protein and evaluated the ability of each peptide to activate HV1s in vitro. Although HV1 T cells did not recognize BCR-ABL fusion epitopes, we found 2 overlapping peptides in the C-terminus of human ABL that strongly activated HV1s (**Figure 3E, F**). Importantly, although the differences in amino acid residues between human and murine ABL at this location were minimal, the corresponding fully murine peptides were completely unable to activate HV1s (not shown). The neoantigenic region therefore resembles an acquired type of mutation akin to resistance mutations, rather than a completely foreign antigen (e.g. OVA). Thus, we generated a physiologically relevant TCR knock-in mouse that produces a population of CD4+ T cells specific for a leukemia neoantigen.

**Figure 3:**
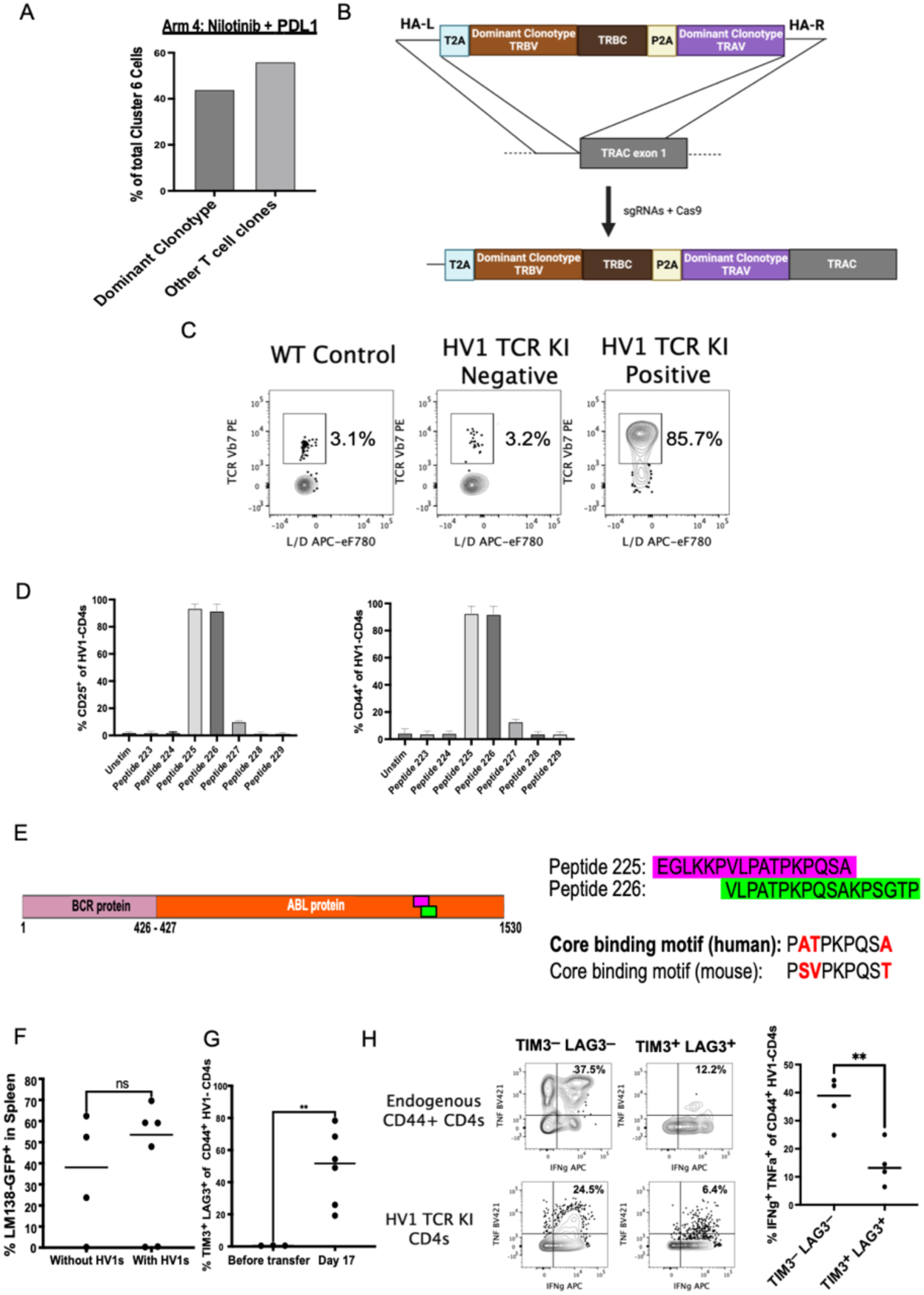
HV1 TCR knock-in mice produce leukemia-specific CD4+ T cells that resemble endogenous anti-leukemia CD4+ T cells. A) Bar graph showing the expansion of the dominant clone within the Tr1-like cluster (Cluster 6) that was identified by scRNA-sequencing of bulk mouse CD4+ T cells from leukemic mice treated with nilotinib +/− anti-PDL1. B) An outline of how the HV1 TCR KI mice were made. The TCR associated with the dominant clone was knocked-in upstream of exon 1 of the *Trac* locus. See Supplementary Methods for more information. C) Representative plots showing the expression of the TCRβ chain associated with the dominant clone in CD4+ T cells from blood of mice that were positive for the HV1 TCR KI vs mice that were negative for the HV1 TCR KI, with WT CD45.2 mice as a control. D) CD4+ T cells were enriched from HV1 TCR KI mice and cultured with T-cell-depleted splenocytes and individual peptides from a peptide library spanning the human BCR-ABL protein. Bar graphs showing the two peptides in the C-terminus of human ABL that induced the upregulation of activation markers CD25 and CD44 on the HV1 T cells. The data is shown as median. E) The core binding motif of the peptide specific for the HV1 TCR was determined using NetMHCIIPan V4.1. The core peptide sequence in the murine vs human version of the peptide is shown, with the differences in amino acid residues highlighted in red. F) Thy1.1 mice were injected with 200,000 Thy1.2+ HV1 T cells, followed by a challenge with 2500 LM138 leukemia cells. Plots showing the leukemia burden in Thy1.1 mice that either did or did not get injected with the HV1 T cells at day 17 post-leukemia challenge. G) Plots showing the proportion of HV1 CD4+ T cells prior to or after transfer that co-express the inhibitory receptors TIM3 and LAG3. H) Plots showing the proportion of IFNγ+ TNF+ cells among TIM3+ LAG3+ vs TIM3-LAG3-HV1 CD4+ T cells. The data is shown as median, with statistical significance determined using an unpaired t test. The data is a summary of 2 independent repeats.

To validate whether the HV1 T cells in leukemic mice recapitulate the Tr1 phenotype we observed in polyclonal CD4+ T cells, we adoptively transferred HV1s into congenically distinct recipients, followed by leukemia challenge. The transfer of 200,000 HV1s significantly prolonged the survival of mice challenged with LM138 leukemia cells and treated with a non-curative course of nilotinib (**Figure 3G**). HV1s from terminally leukemic mice upregulated TIM3 along with multiple other inhibitory receptors including LAG3. Furthermore, TIM3+/LAG3+ HV1s had an impaired ability to produce TNF upon ex-vivo stimulation (**Figure 3H**), although IFNγ production remained robust. Thus, HV1 T cells have transient anti-leukemia activity and recapitulate the phenotype and functional state of polyclonal CD4+ T cells during leukemia challenge.

### The leukemia microenvironment superimposes a Tr1 terminal fate onto canonical CD4+ T-cell subsets

During leukemia progression, HV1s progressively downregulated the key helper cytokine, CD40L, while upregulating IL10 upon ex vivo stimulation (**Figure 4A, B**). Consistent with this, HV1s from day 14 leukemic mice suppressed bystander T cells ex vivo (**Figure 4C-F**). Both IL10-GFP+ and IL10-GFP-HV1s suppressed bystander T cells, suggesting that the suppressive function of HV1s is not solely dependent on IL10 production (**Figure 4E, F**). Thus, leukemia programs HV1 cells in vivo towards a phenotypic Tr1 state. Next, we examined whether this state was distinct from T-cell exhaustion. Chronically stimulated CD8+ T cells experience progressive epigenetic silencing of effector cytokines, which is a definitive feature of terminal exhaustion^20,21^. To determine whether this occurs in leukemia-specific CD4+ T cells, we characterized the epigenetic state of HV1 cells during the course of leukemia progression using scATAC-seq. We transferred HV1 T cells into congenically distinct hosts. One day later, we transferred 2500 LM138 cells. HV1 T cells were FACs-sorted 9, 13, and 17 days later. As a control, naïve HV1 cells were also sorted from TCR knock-in mice (**Figure 5A**). Despite the fact that HV1 T cells express a monoclonal TCR, we observed heterogenous differentiation of HV1s, which continued to dynamically change even at late timepoints (**Figure 5B, C**). By day 9 virtually all transferred HV1 T cells had encountered antigen and became activated. Day 9 HV1 cells were dominated by an early-activation signature (cluster 4) and a Th1-like signature (cluster 1), on the basis of *Pdcd1* and *Ifng* accessibility. By day 13, early-activated T cells had increased *Cxcr5* and *Il21* accessibility, suggestive of a T follicular helper (Tfh) identity^22,23^. This bifurcation into a Tfh or T-helper state resembles canonical differentiation of naïve CD4+ T cells in response to infectious pathogens^24–26^. Thus, HV1s appear to follow a conserved differentiation pathway at early timepoints after leukemia challenge.

**Figure 4:**
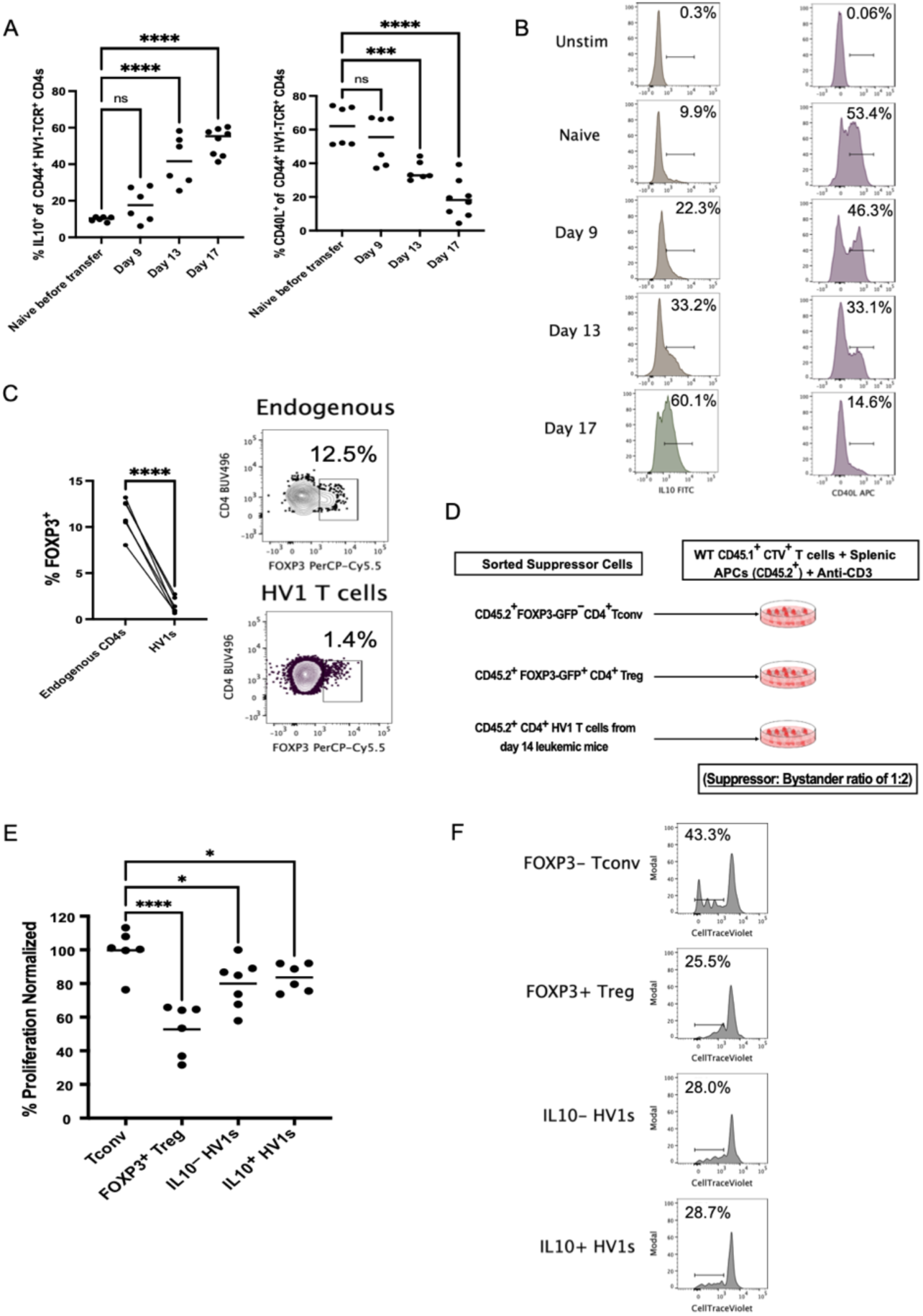
HV1 TCR knock-in CD4+ T cells recapitulate the leukemia-induced Tr1 phenotype in vivo. Thy1.1 mice were injected with 200,000 Thy1.2^+^ HV1 T cells, followed by a challenge with 2500 LM138 leukemia cells. A,B) Graphs and representative flow plots showing the proportion of IL10+ and CD40L+ cells among ex vivo stimulated HV1 T cells from leukemic mice at the indicated time points. C) Graph and representative plot showing the proportion of FOXP3+ cells among endogenous Thy1.1^+^ and transferred Thy1.2^+^ HV1 CD4+ T cells from day 13 leukemic mice. The results are a summary of 2 experiments. CD45.1+ mice were injected with 100,000 CD45.2+ HV1 T cells, followed by a challenge with 2500 LM138 leukemia cells. D) IL10-GFP+ and IL10-GFP − HV1s were sorted from Day 14 leukemic mice and cultured with bystander CTV+ T cells with APCs and anti-CD3. E,F) Graphs and representative flow plots showing the proportion of bystander T cells that proliferated. The results are a summary of 2 experiments. Statistical significance was determined using a one-way ANOVA, followed by the Holm-Sidak’s multiple comparisons test. *p<0.05, ***p<0.001, ****p<0.0001.

**Figure 5:**
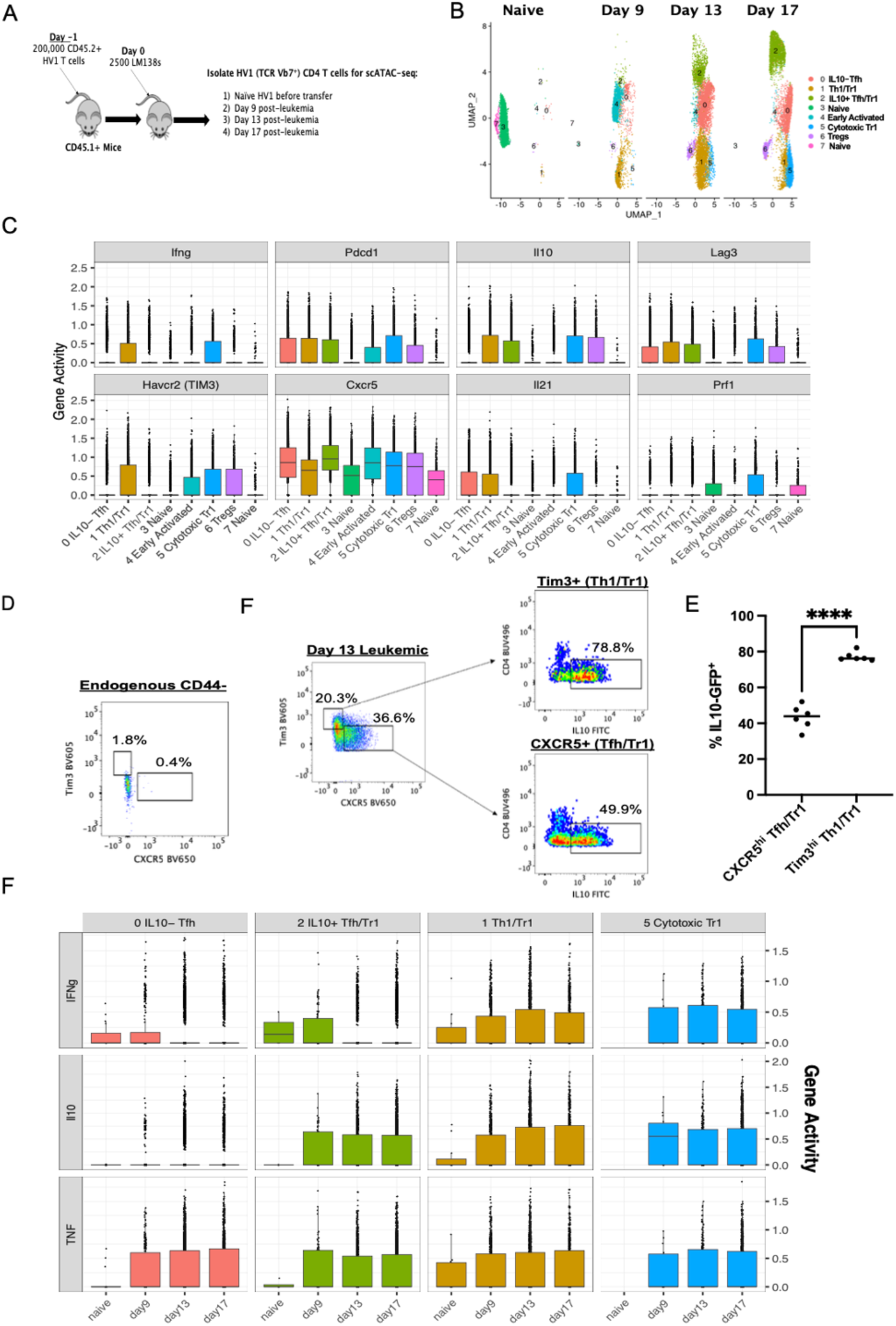
Both Th1 and Tfh-like CD4+ cells acquire an IL10+ Tr1-phenotype that is distinct from exhausted CD8+ T cells. A) Schematic describing the scATAC-seq experiment of HV1 T cells from leukemic CD45.1+ mice at different time points post-leukemia challenge. B) UMAP plot showing the different clusters of HV1 T cells identified at each time point. The number of cells at each time point was as follows: naïve = 3488, Day 9 = 2912, Day 13 = 7897, Day 17 = 7471. C) Box plots showing the gene activity score (readout for accessibility) of the indicated genes in the different CD4+ T-cell clusters. D,E) Representative flow plots and graph showing the proportion of IL10-producing cells within ex vivo stimulated HV1s from day 13 leukemic mice, stratified by expression of Cxcr5+ (Tfh) or Tim3+ (Th1). The results are a summary of 2 experiments. Statistical significance was determined using an unpaired t test. ****p<0.0001. F) Box plots showing the gene activity score for the indicated cytokine-associated genes in the indicated clusters of HV1s T cells at the different time points post-leukemia challenge. Also see Figure S3, S4.

During later timepoints, HV1s continued to change. Both Th1 and Tfh cells developed accumulative epigenetic changes consistent with progressive adoption of a Tr1 identity. For example, by day 9, Th1-like cells (cluster 1) had accessible loci in genes associated with Tr1 identify including *Il10* and *Lag3*. Likewise, by day 13, a Tfh-like cluster (cluster 2) had also upregulated *Il10* and *Lag3* (**Figure 5D, E, Supplementary Figure 2**). Finally, day 17 HV1 T cells largely consolidated into three groups: an IL10+ Tfh/Tr1 state (cluster 2), an IL10-Tfh state (cluster 0) and an IL10+ Tr1/cytotoxic state (Clusters 1/5). Moreover, while the IL10+ Tfh/Tr1 subset exhibited many characteristics of classical Tfh cells, they failed to maintain accessibility of the key cytokine gene *Il21*. While terminally-differentiated Tfhs are known to lose IL21 production, they typically maintain locus accessibility within the *Il21* gene^27^; therefore, our findings suggest that leukemia superimposes an epigenetically distinct Tr1-like state onto early Tfh cells. Finally, although FOXP3+ Tregs did emerge upon encounter with leukemia, they represented a small subset (cluster 7). Importantly, Th1/Tr1 cells upregulated *Havcr2* but showed no evidence of losing accessible loci within *Ifng* or *Tnf* during the course of leukemia progression, suggesting a functional state distinct from terminal exhaustion (**Figure 5F**).

### IL10R and PDL1 blockade differentially impact T-cell differentiation and leukemia relapse

We previously showed that leukemic mice treated with the BCR-ABL inhibitor, nilotinib, experience a transient remission followed by nearly universal relapse. In contrast, adding PDL1 blockade to nilotinib cured 70% of leukemic mice^5^. As leukemia-specific CD4+ T cells increasingly acquire a Tr1 phenotype during leukemia progression, we decided to determine the contribution of leukemia-specific Tr1s towards leukemia relapse. Our data and data from other groups suggest that Tr1s can mediate their suppressive activity in an IL10-independent manner^28^. However, T-cell-intrinsic IL10/IL10R signaling has been shown to be essential for maintaining a Tr1-phenotype in vivo. We thus evaluated the ability of IL10R blockade to prevent relapse in our leukemic mice in the context of combined treatment with nilotinib. Mice were injected with 10,000 HV1 T cells and then received 2500 LM138 leukemia cells 24 hours later. Treatment with isotype control antibody, nilotinib alone, nilotinib plus anti-IL10R antibody, or nilotinib plus anti-PDL1 antibody was begun on day 14 after leukemia transfer. While combined PDL1 blockade and nilotinib treatment reduced leukemia growth (**Supplementary Figure 3**) and prolonged survival, combined IL10R blockade and nilotinib had no impact on leukemia burden or survival (**Supplementary Figure 3**).

We next performed scRNA-seq on HV1 CD4+ T cells and CD44+ CD8+ T cells from untreated leukemic mice as well as from leukemic mice treated with nilotinib + isotype, IL10R, or PDL1 blocking antibodies (**Figure 6A**). The HV1 CD4+ T-cell clusters from the scRNA-seq data appeared to approximate the populations of CD4+ T cells we identified by scATAC-seq (**Figure 6B, C**). Th1-like cells (clusters 1 and 2) were distinguishable from Tfh-like cells (cluster 3) based on the mutually-exclusive expression of *Blimp1* and *Bcl6*, respectively. Cluster 1 had increased expression of *Il10, Maf* and *Havcr2*, consistent with a TIM3+ Tr1 population that emerges early in leukemia progression from Th1s. Although cluster 0 did express Tfh-associated genes, including *Il21 and Cxcr5*, it had lower expression of *Bcl6* and higher expression of *Tcf7* and *Lef1,* known markers of stem-like T cells (**Figure 6C**). In contrast, cluster 3 was enriched in genes associated with terminally-differentiated T cells such as *Crtam, Eomes,* and *Slamf7* (**Figure 6C**). Thus, cluster 3 is likely an “effector” Tfh population while cluster 0 is likely a stem-like population with the potential for Tfh differentiation. Cluster 5 cells overexpressed the activation markers *Nr4a1-3*, *Nfatc1,* as well as *Myc*. Cluster 6 and 7 were small populations that expressed markers associated with proliferation (*Ki67*) and Tregs (*Foxp3*, *Il2ra*) respectively (**Figure 6C**). We also used label transfer-based approaches, which further demonstrated concordance between clusters derived from scATAC-seq and scRNA-seq datasets (**Supplementary Figure 4**).

**Figure 6:**
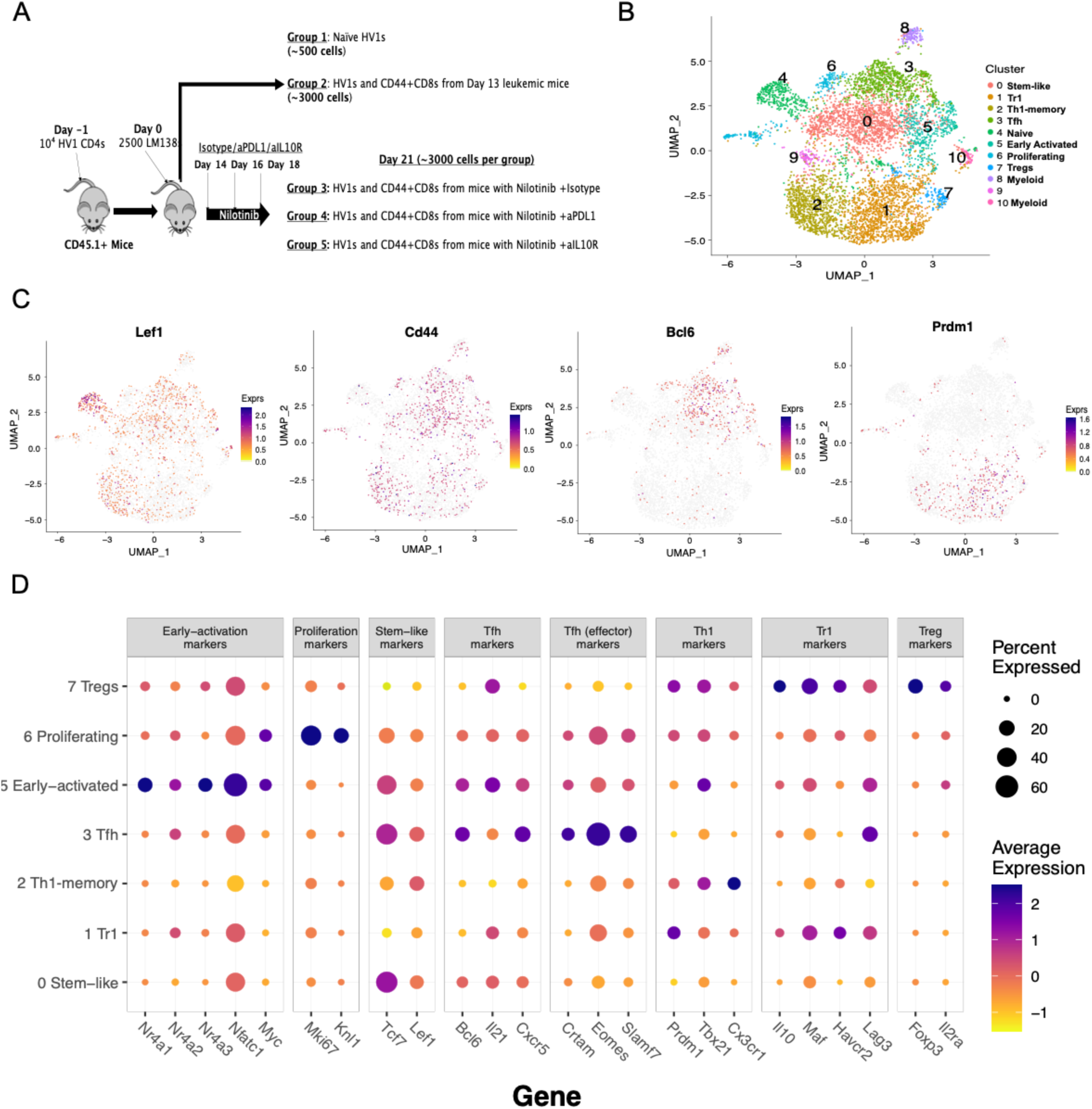
scRNA-seq recapitulates key leukemia-specific CD4+ T-cell subsets from mice with B-ALL. A) Schematic describing the scRNA-seq experiment of HV1 T cells from leukemic CD45.1+ mice in the different treatment arms. B) UMAP plot showing the different clusters of HV1 T cells identified. C,D) Feature plots and dot plot showing the expression of the indicated genes within the different CD4+ T-cell clusters. (Also see Figure S4).

**Figure 7:**
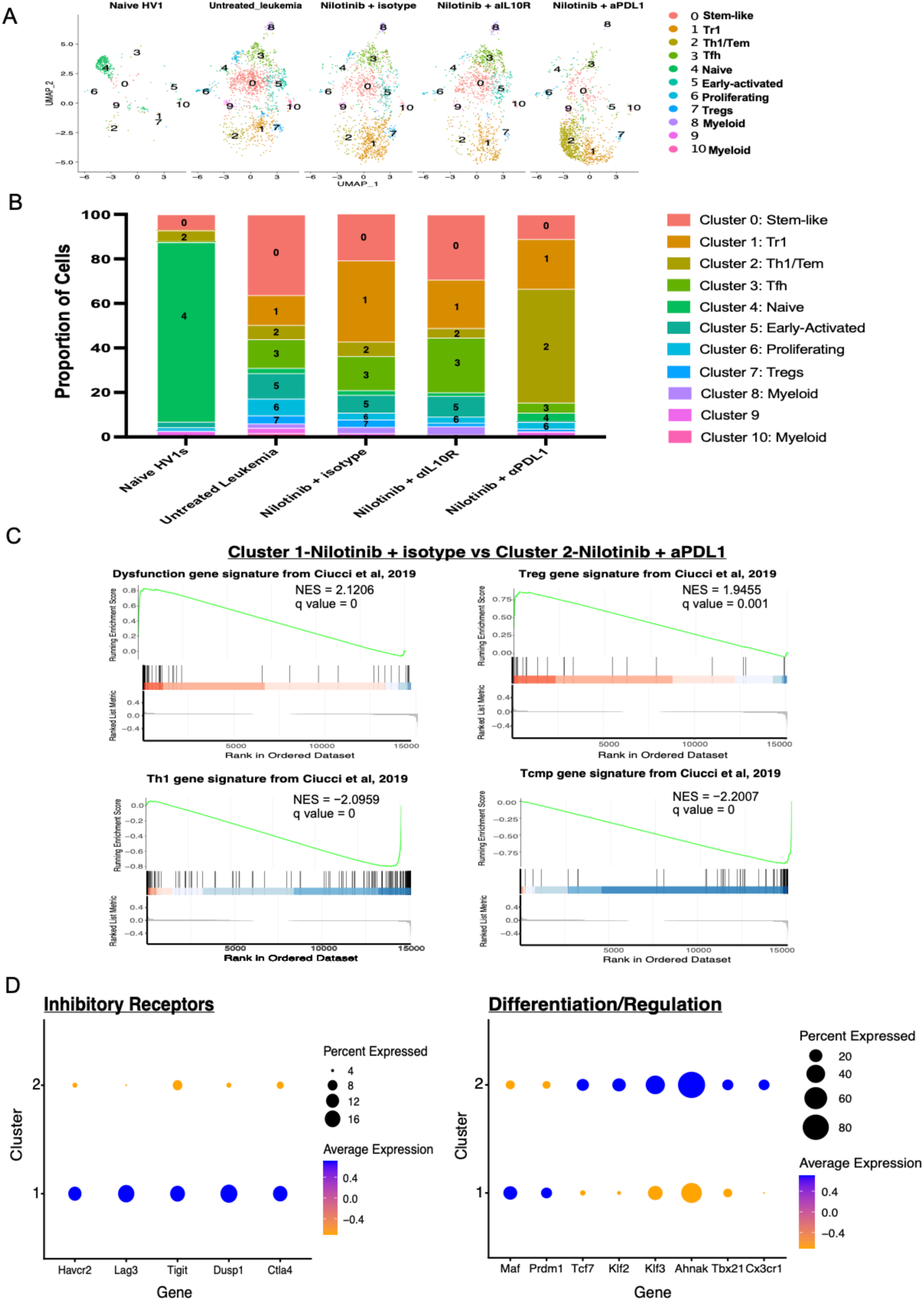
IL10R blockade depletes Tr1s, while PDL1 blockade directs CD4+ T cells towards a Th1 fate. A) UMAP plot showing the different HV1 CD4+ T-cell clusters stratified by treatment arm. The number of cells at each time point was as follows: naïve HV1 = 378, untreated leukemia = 1561, nilotinib = 1407, nilotinib + anti-IL10R = 1198, nilotinib + anti-PDL1 = 1805. B) Bar plot showing the proportion of cells within the different clusters that were present in HV1 T cells from each treatment arm. C) GSEA plots showing enrichment of key gene signatures that differentiate cluster 1 and cluster 2 cells from mice treated with nilotinib +/− anti-PDL1. D) Dot plots showing the expression of the indicated genes associated with inhibitory receptors and differentiation/regulation within cluster 1 and 2. (Also see Figure S5)

To contrast how IL10R and PDL1 blockade impacted CD4+ T-cell differentiation, we subsetted the HV1 CD4+ T-cell clusters by treatment arm (**Figure 6A, B**). Mice treated with nilotinib alone had a further increase in Tr1s (cluster 1) and a reduction in stem-like cells (cluster 0) compared to untreated leukemic mice prior to nilotinib treatment. As expected, mice treated with nilotinib and IL10R blockade had a lower proportion of Tr1s; this suggests that IL10R blockade either prevented the expansion of pre-existing Tr1s or the differentiation of stem-like cells into a Tr1 state. In contrast, mice treated with nilotinib and PDL1 blockade not only had a lower proportion of Tr1s (cluster 1) but they also had a significant expansion of Th1s (cluster 2, **Figure 6B**). Comparison of DEGs between cluster 1 and cluster 2 revealed that cluster 1 cells overexpressed *Hav*c*r2, Lag3, Maf,* and *Prdm1* (**Figure 6C, D**). In contrast, cluster 2 had higher expression of *Tbx21, Cx3cr1,* and *Tcf7* (**Figure 6C, D**). The top differentially-expressed gene in cluster 2 vs cluster 1 was *Ahnak*, which was also found to be relatively overexpressed in Th1s relative to Tr1s in clinical samples (**Figure 1B**). Thus, nilotinib and PDL1 blockade not only reduce Tr1s (cluster 1) but also expand Th1s (cluster 2). Notably, both Th1 and Tr1 cells equally expressed markers of cytotoxicity including *Nkg7* and *GzmB*.

We next examined CD8+ T-cell subsets from the various treatment arms to understand how PDL1 and IL10R blockade impacted antigen-experienced (CD44+) CD8+ T cells. We had previously shown that the frequency of cytotoxic (GzmB+) CD8+ T cells increased after treatment with nilotinib + anti-PDL1 therapy. We now observed that in untreated leukemia-bearing mice, nearly all CD8+ T cells belonged to a single cluster with *Tcf7* and *Bach2* overexpression, consistent with a naïve or early central-memory phenotype (cluster 0, **Supplementary Figure 5**A**, B**). This subset bore few hallmarks of recent activation even at high leukemia burdens, suggesting that the majority of antigen-experienced CD8+ T cells do not directly participate in leukemia clearance. Yet, after treatment with nilotinib, we observed emergence of a population (cluster 1) of intermediate effectors based on the upregulation of *Tox, Tbx21, Havcr2,* and downregulation of *Tcf7* (**Supplementary Figure 5**B**, C**). In contrast to CD4+ T cells, the addition of anti-PDL1 or IL10R blockade only modestly modified the distribution of CD8+ T-cell subsets (**Supplementary Figure 5**B). CD8+ T cells also continued to express *Ifng, Prf1,* and *GzmB,* arguing against a state of terminal exhaustion (**Supplementary Figure 5**C). Analysis of CD8+ T-cell clusters from clinical specimens also revealed analogous subsets to those observed in our mouse models (**Supplementary Figure 6**); here, we also observed stem-like CD8+ T cells (cluster 0) that co-expressed *Tcf7* and *Tox* and the effector cytokines, *Ifng and Tnf*. We also saw a subset of effector-memory CD8+ T cells (cluster 4) that overexpressed *Klrg1, Tbx21, Tcf7, and Il7R.* This subset also overexpressed cytotoxicity-associated molecules (*Prf1, Gzmb*) but lacked expression of effector cytokines, consistent with a known switch from cytokine production to cytotoxicity. We also saw CD8+ T cells (cluster 8) that overexpressed cytokine and cytotoxicity-associated genes as well as multiple inhibitory receptors. This subset had an intermediate level of *Tbx21* expression, with minimal upregulation of *Tox,* resembling a population of “intermediate effector” T cells that have not reached terminal exhaustion^29^. Thus, CD8+ T cells largely harbor transcriptional states consistent with naïve, stem-memory, or early-activated effector subsets. Curative therapy with nilotinib + anti-PDL1 correlates with a polarization of CD4+ T cells towards a CD40L^hi^/IL10^lo^ Th1 state and improved recruitment of cytotoxic CD8+ T cells but with minimal effect on CD8 transcriptional states. The further supports a mechanism whereby anti-PDL1 blockade primarily functions via modulation of CD4+ T-cell helper function and CD8+ recruitment, rather than direct effects on CD8+ T-cell function.

## Discussion

Our findings indicate that neoantigen-specific CD4+ T cells adopt Tr1 tolerogenic fates in the leukemia microenvironment and thus inhibit the activation of cytotoxic T cells. Repolarization of CD4+ T cells towards Th1 function correlates with improved CD8+ T-cell recruitment and reduction or elimination of residual leukemia.

Our studies using the HV1 knock-in mouse allowed us to perform an in depth analysis of the differentiation pathways and terminal fates of neoantigen-specific CD4+ T cells during ALL progression. HV1s developed along a conserved bifurcation into Tfh and Th1 subtypes shortly after initial priming. However, by late timepoints, both Tfh and Th1 precursors largely adopted FOXP3-/IL10+ states and suppressive functions, consistent with a Tr1 fate. We observed that gene expression patterns of leukemia-specific Tr1s is similar to tolerogenic CD4+ T cells induced by the normal hematopoietic stem cell niche^11^. This suggests that acute leukemia may co-opt signals of a pre-existing immune circuit between HSCs and CD4+ T cells as a mechanism of escaping immune pressure. While we focused on the significance of Tr1s towards relapse in B-ALL, it is plausible that Tr1s may also broadly contribute to leukemia initiation by suppressing a cytotoxic response towards a variety of mutation-harboring progenitor cells. This may represent a requisite step in the development of conditions ranging from clonal hematopoiesis of indeterminate origin to myelodysplastic syndrome and acute leukemias.

A variety of Tr1-mediated suppressor functions in infectious, transplant, and neoplastic settings have been well described previously, and center on IL10-dependent and -independent mechanisms^30–32^. While serum IL10 level alterations are associated with the pathogenesis of B-ALL^33^, we found little evidence that neutralization of IL10 signaling improves leukemia control. As expected, we found that Tr1s can directly suppress CD8+ T cells in ex-vivo co-culture assays, but CD8+ T cells from in vivo models or clinical samples showed minimal intrinsic functional deficits. Instead, they appear to harbor signs of minimal activation, as reflected by a predominantly naïve or early stem cell memory transcriptional state. Tr1 cells may contribute to inadequate CD8 cross-activation by direct lysing MHC-II+ tumor antigen-presenting type 1 conventional dendritic cells^34^. Consistent with this, we observed a robust cytotoxic signature in Tr1s in both mouse models and clinical specimens.

Clinical outcomes among patients with B-ALL have markedly improved due in part to the advent of T-cell mediated immune therapies^35,36^. However, relapse after T-cell therapies remains common, emphasizing the capacity of residual leukemia cells to escape immune pressure. CD4+ T cells serve a critical role in hematologic malignancies by integrating recognition of MHCII-restricted neoantigens with a downstream effector or suppressive response. Consistent with this, phenotypically dysfunctional TIM3+ CD4+ T cells are highly predictive of relapse in B-ALL^3,4^. As our studies show that TIM3 expression is largely confined to Tr1 cells in both mouse models and clinical samples, we suggest that relapse-associated TIM3+ CD4+ T cells are likely to be predominantly comprised of Tr1s. This will require confirmation in a larger cohort of patients. However, TIM3+ CD4+ T cells are already known to be associated with relapse after allogeneic HSC transplant and resistance to the T-cell engager, blinatumomab^37,38^. Therefore, Tr1 cells may convey resistance not only to endogenous immune surveillance but also towards contemporary T-cell bispecific engagers.

Critically, we suggest that the suppressive effects of Tr1s can be overcome by rebalancing the local population of neoantigen-specific CD4+ T cells towards CD40L^hi^ pro-helper states. We predict this occurs via restoration of 3-cell immune structures called triads, formed between Th1, cDC1, and CD8+ T cells, that lead to robust CD8 T-cell cross-activation^39^. Efforts to accomplish this therapeutically may synergize with existing T-cell therapies by establishing an environment that is permissive for cytotoxic attack towards residual leukemia cells^40^.

## Acknowledgments

The authors thank G. Hubbard for assistance with mouse procedures; Shelley, T. Martin, J. Motl, and P. Champoux for cell sorting and maintenance of the Flow Cytometry Core Facility at the University of Minnesota (5P01AI035296); W. Lahr, Y. You, the University of Minnesota Genome Engineering Shared Resource (GESR), and the Mouse Genomics Lab (MGL) for helping make the HV1 TCR KI mouse; M.R. Rollins and I.M Stromnes for guidance in making and validating the HV1 TCR KI mice; and P. Grady and UMGC core for assistance with scRNA-seq and scATAC-seq. The authors also acknowledge the Minnesota Supercomputing Institute (MSI) at the University of Minnesota for providing computational resources, and the Hematologic Malignancies Tissue Bank for providing clinical samples. This project was supported by funding from the Children’s Cancer Research Fund (SIT and MAF), funding from the Department of Laboratory Medicine and Pathology (MAF), and NIH grant R01 AI124512 (MAF).

## Authorship Contributions

H). V., S.I.T and M.A.F designed experiments; H.V., S.I.T, L.H.H and M.A. performed experiments and analyzed data; T.P.K., A.H. and Y.Q. analyzed data; B.R.W. helped design the adenoviral vectors for the HV1 TCR KI mouse model; and H.V., S.I.T, and M.A.F. wrote the manuscript.

## SUPPLEMENTAL FIGURES

**Supplementary figure 1:**
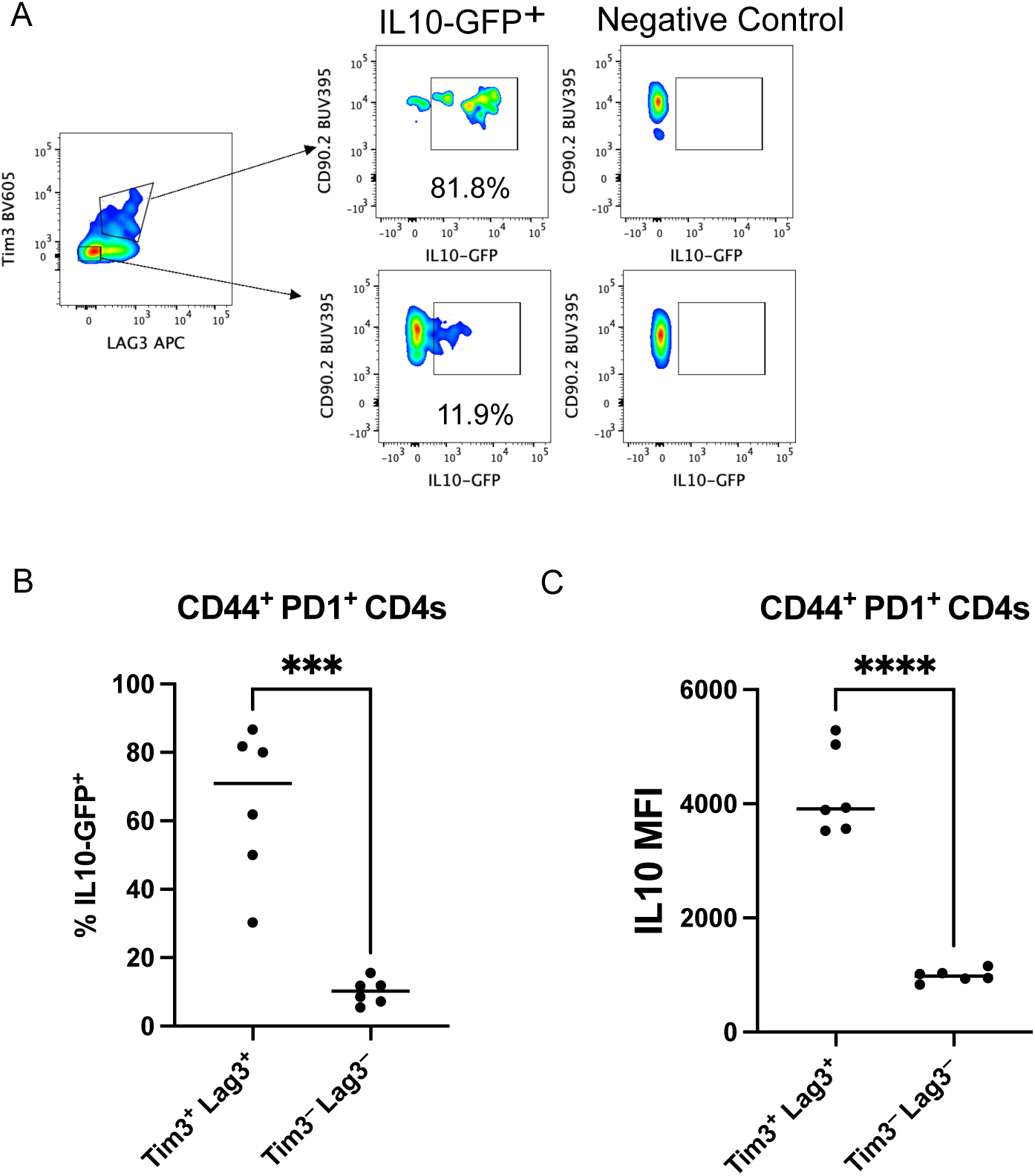
CD4 T cells with a Tr1-like phenotype are present in Pax5^+/−^ x EBF^+/−^ leukemias. WT IL10-GFP^+^ mice were challenged with 5 × 10^5^ Pax5^+/−^ x EBF^+/−^ leukemia cells. A) Representative flow plots and B) Plots showing the proportion of IL10-GFP^+^ T cells among Tim3^+^ Lag3^+^ vs Tim3^−^ Lag3^−^ CD4 T cells (n=6 mice per group). The graph is a summary of 2 independent experiments shown as a median, with statistical significance determined using an unpaired t-test.

**Supplementary figure 2:**
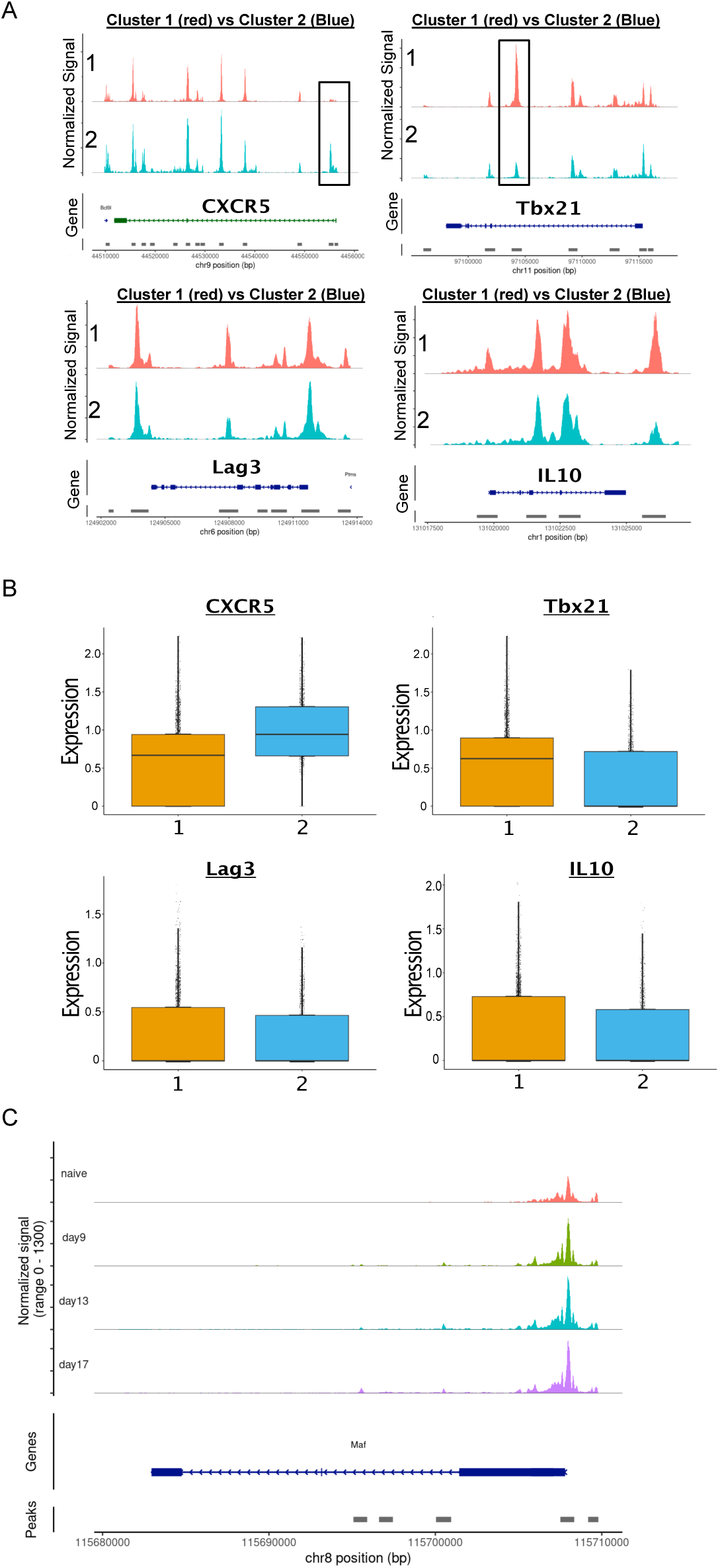
Leukemia cells induce an epigenetic Tr1-like program over time in Th1 and Tfh-polarized HV1 CD4 T cells. Accessibility traces (A) and predicted expression (B) of key Th1/Tfh genes (Tbx21, CXCR5) and key Tr1 genes (IL10, Lag3) within cluster 1 vs cluster 2 of HV1 CD4 T cells at Day 13 post-leukemia challenge. C) Accessibility trace of c-Maf, a key Tr1-associated transcription factor, in HV1 T cells at the indicated timepoints post-leukemia challenge

**Supplementary figure 3:**
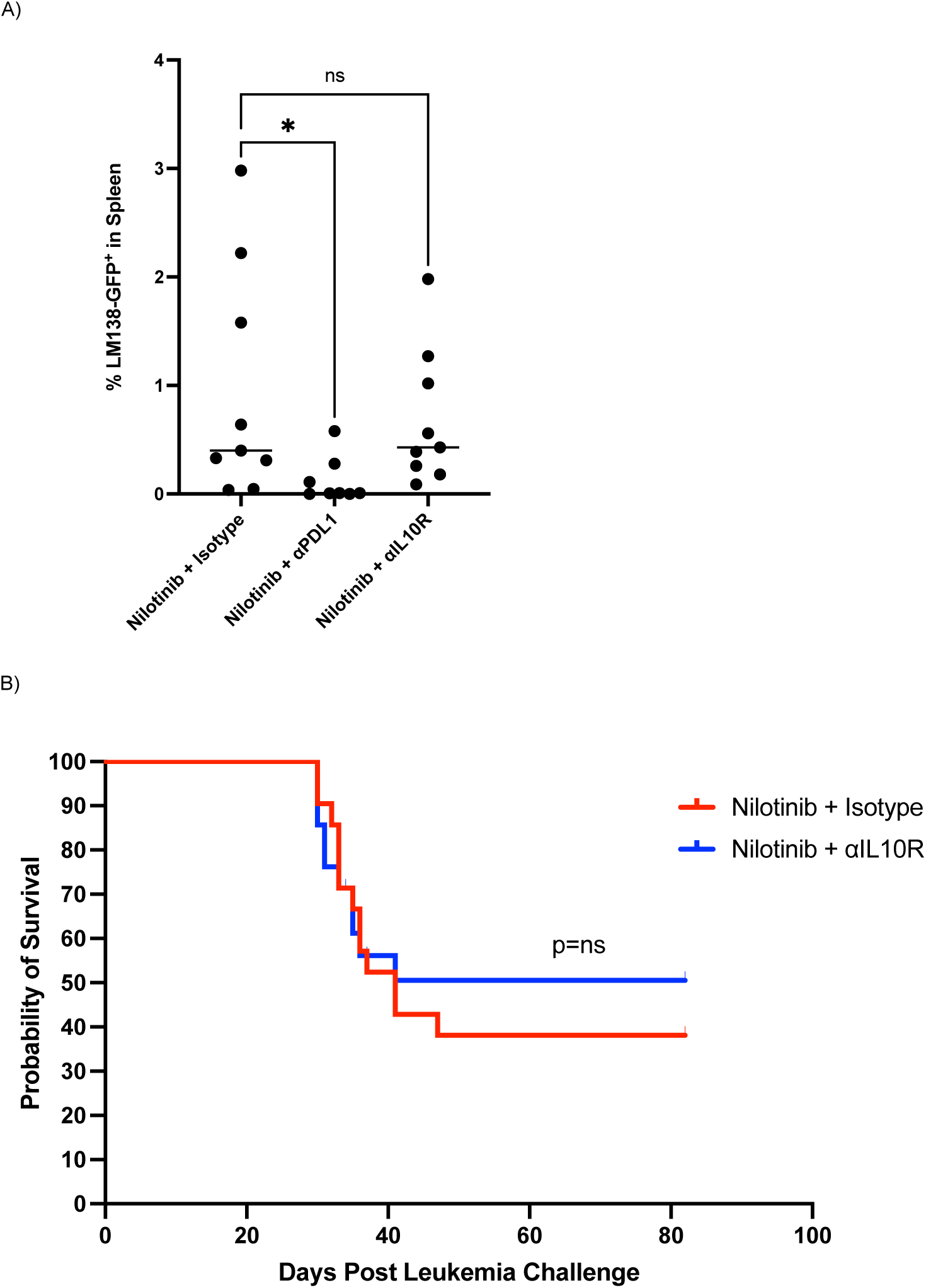
PDL1 blockade, but not IL10R blockade, inhibits leukemia growth and prevents relapse in the context of nilotinib treatment. A) 10,000 HV1 T cells were transferred into CD45.1 mice 1 day prior to challenge with LM138 leukemia cells, followed by treatment with nilotinib + isotype, IL10R blocking or PDL1 blocking antibodies at day 14 post-leukemia challenge (See methods for treatment details). The leukemia burden in the spleen of the mice at day 21 post-leukemia challenge is shown. The plot is a summary of 2 independent experiments as a median with n=4 mice per group. Statistical significance was analyzed using a one-way ANOVA test. *: p<0.05 B) Kaplan-Meier curves showing the survival of CD45.1 mice where 10,000 HV1 T cells were transferred 1 day prior to challenge with LM138 leukemia cells, followed by treatment with nilotinib + isotype or IL10R blocking antibodies at day 14 post-leukemia challenge (See methods for treatment details). The plot is a summary of 2 independent experiments with n=10 mice per group. Statistical significance was analyzed using the Mantel-Cox log-rank test.

**Supplementary figure 4:**
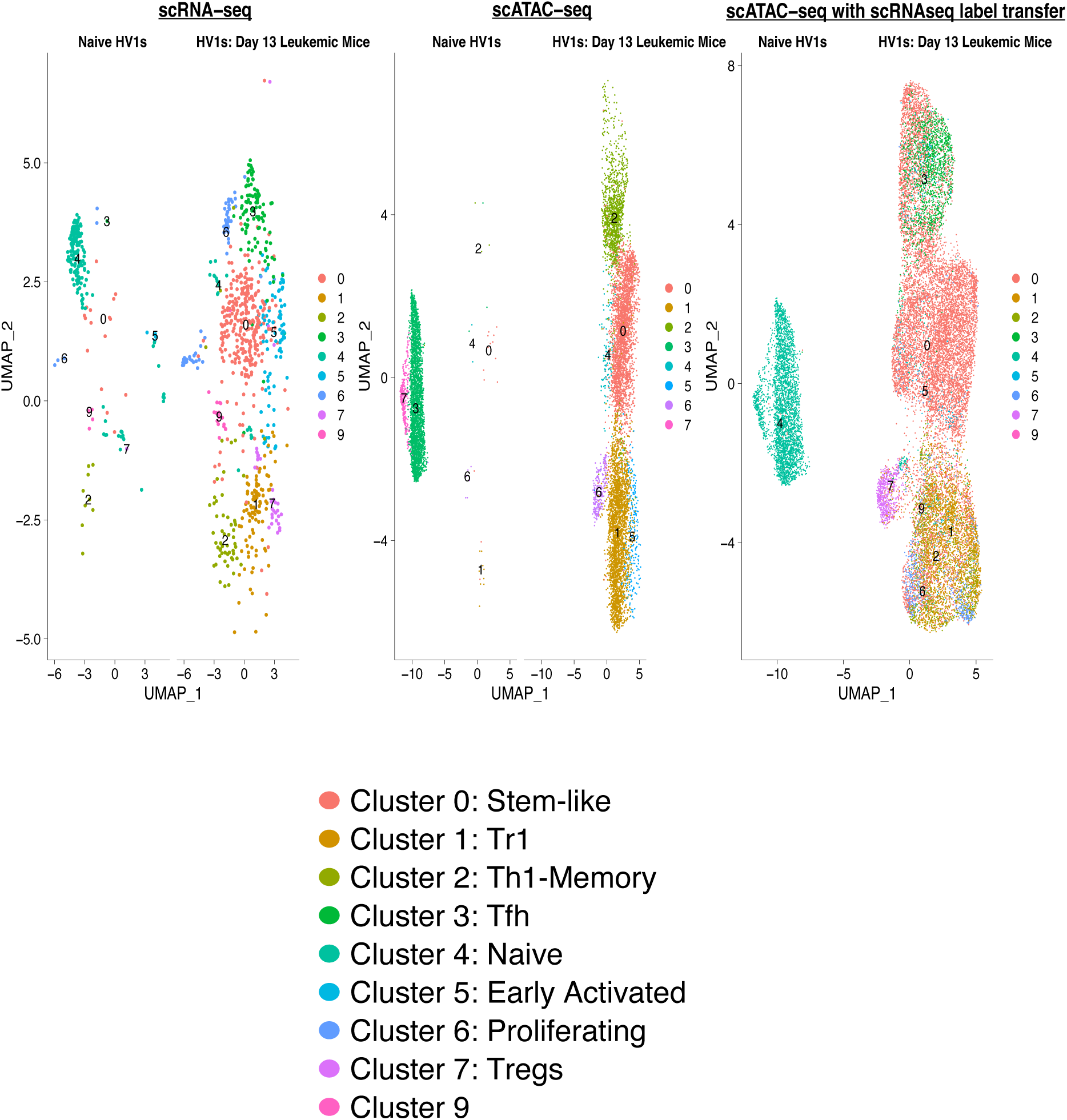
Label-transfer approaches. UMAP plots showing HV1 CD4 T cells after integration of scRNA-seq and scATAC-seq data from naïve TCR transgenic mice or untreated leukemic mice.

**Supplementary figure 5:**
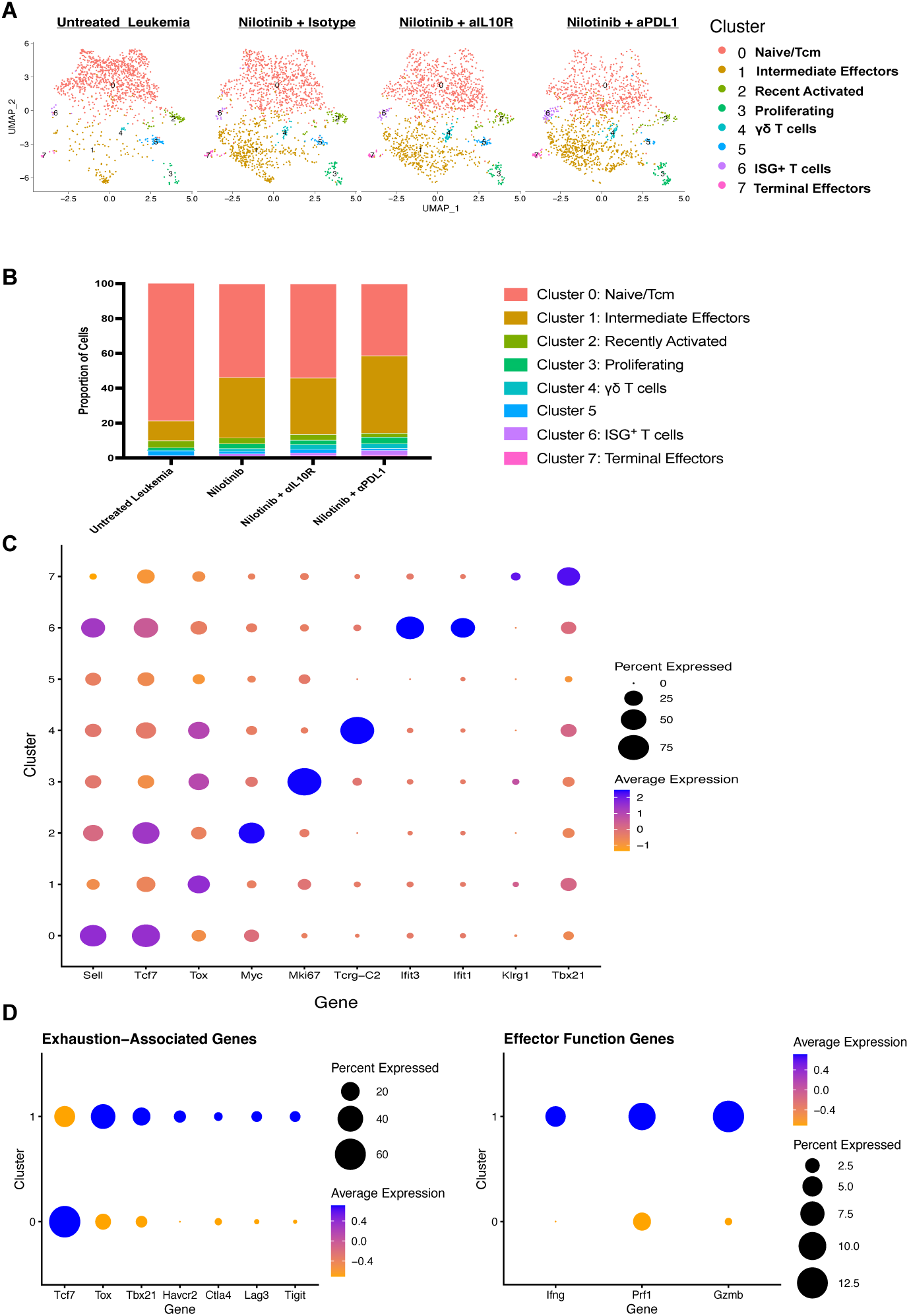
Analysis of CD8^+^ T cells from murine scRNA-seq analysis shows that leukemia impairs CD8^+^ T cell activation, which is enhanced by nilotinib and PDL1 blockade. A) UMAP plot showing different CD8**^+^** T cell clusters from leukemic mice in the scRNA-seq dataset (see figure 4 for outline of scRNA-seq experiment). B) Plot showing the proportion of the different CD8**^+^** T cell clusters in the leukemic mice in the different treatment arms C) Dot plot showing the relative expression of key genes that distinguish the different clusters D) Dot plots highlighting the expression of genes associated with exhaustion and effector function in CD8**^+^** T cells from cluster 0 vs cluster 1-the most dominant CD8**^+^** T cell clusters in the dataset.

**Supplementary Figure 6:**
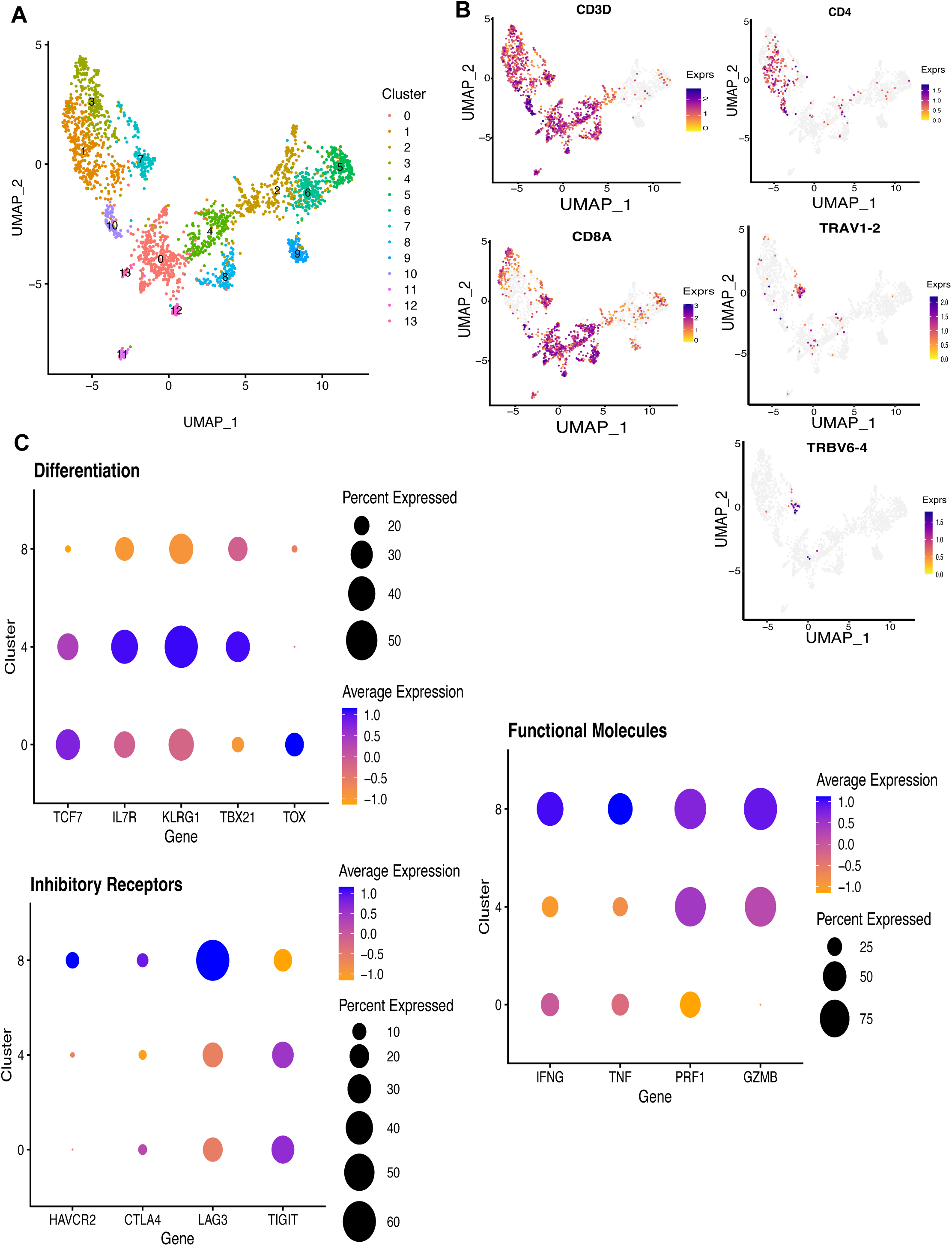
CD8^+^ T cells from clinical specimens show minimal evidence of exhaustion. A) UMAP plot showing different CD8^+^ T cell clusters from a previously-published scRNA-seq dataset of bone marrow biopsies of 5 B-ALL patients at diagnosis^1^. B) Feature plots showing the expression of key genes associated with conventional T cells (CD3D, CD4 and CD8), as well as MAIT cells (TRAV1-2, TRBV6-4). C) Dot plots highlighting the expression of genes associated with differentiation, inhibitory receptors, and effector function in the different CD8^+^ T cell clusters in the dataset.

